# FibroTrack: A Standalone Deep Learning Platform for Automated Fibrosis Quantification in Muscle and Cardiac Histology

**DOI:** 10.1101/2025.09.02.673720

**Authors:** Anas Odeh, Rahaf Salem, Maher Abu Saleh, Ariel Shemesh, Polina Stein, Ido Livneh, Peleg Hasson

## Abstract

Accurate fibrosis quantification is essential for understanding muscle and cardiac disease, yet current manual and semi–automated methods remain slow, subjective, and poorly reproducible. We introduce FibroTrack, a standalone deep learning platform with a graphical user interface (GUI) that fully automates fibrosis analysis across Sirius Red (SR), Masson’s Trichrome (MT), and immunohistochemistry (IHC) stainings. FibroTrack uniquely integrates LAB (lightness, green–red, blue–yellow) color space normalization with a You Only Look Once version 11 (YOLOv11) segmentation model trained on 2,034 histological images. This approach achieved >97% precision and demonstrated excellent concordance with blinded pathologists (Spearman correlation, r = 0.87–0.96). Automated outputs include segmented images and structured spreadsheets, ensuring high reproducibility and scalability. By combining advanced color analysis with state–of–the–art segmentation in an accessible tool, FibroTrack provides a novel, accurate, and clinically relevant solution for high–throughput fibrosis quantification in both preclinical research and pathology practice.

---

Fibrosis is a key biological process essential for tissue repair following injury. However when dysregulated, it contributes to the pathology of numerous chronic muscle diseases, such as Duchenne Muscular Dystrophy (DMD) and cardiovascular disorders such as heart failure with preserves ejection fraction (HFpEF)^1,2^. During the fibrotic process, excessive extracellular matrix (ECM) deposition disrupts normal tissue architecture and function ultimately inhibiting recovery^3^. Reliable quantification of fibrosis is a critical tool in preclinical research and, in specific clinical cases, can provide insights into disease progression, prognosis, and therapeutic response. However, current methodologies for fibrosis assessment, primarily manual and semiquantitative ^4,5^ , are fraught with limitations. These techniques are labor-intensive, prone to observer variability, and lack the precision required for high-throughput analysis, making them insufficient for the rigorous demands of modern fibrotic research.

A key issue for many users is the ease of use including all stages of the process ranging from image(s) input, analysis and output. Here we describe the development of FibroTrack, a deep learning-based graphical user interphase (GUI) designed to automate the quantification of fibrosis in histological images of muscle tissues. This GUI represents a significant advancement in fibrosis analysis by providing a robust, reproducible, and efficient tool capable of processing large datasets of Sirius red (SR), Masson’s Trichrome (MT) and immunohistochemistry (IHC)-stained images. The GUI is capable of accurately segmenting fibrotic and non-fibrotic regions and calculate fibrosis ratios. These fibrosis ratios are expressed as the percentage of fibrotic area relative to total muscle tissue area. The GUI’s ability to generate an easy-to-interpret, structured outputs, such as organized Excel files, offers a marked improvement over traditional methods, ensuring consistent and unbiased measurements across different experimental conditions.

Notably, FibroTrack is standalone, and supports all common image formats, including JPG, PNG, and TIFF. This versatility, coupled with an intuitive user interface that does not require any computing skills renders this GUI highly accessible to a broad range of researchers, from basic scientists to clinicians. Importantly, the GUI’s deep learning model has been rigorously trained and validated on a diverse dataset, ensuring its generalizability across different staining protocols and different muscle types. By streamlining fibrosis quantification, our GUI addresses a major bottleneck in fibrotic disease research, enabling rapid and accurate analysis that is essential for both experimental and translational studies.

Here, we detail the development, validation, and application of FibroTrack, demonstrating its utility in quantifying fibrosis in both skeletal and cardiac muscle tissues. We also demonstrate evidence of its performance in terms of accuracy, reproducibility, and scalability, highlighting its potential as a valuable tool for advancing fibrosis research in both preclinical models and clinical samples. Given the increasing recognition of fibrosis as a therapeutic target ^6^ across multiple disease domains, FibroTrack will be an ideal resource for studies requiring precise and high-throughput fibrosis quantification.

## Results

Although multiple image analysis software or plugins are available, many of them require much work from the user, whether in arranging the images, various segmentation steps or in organizing the detailed output altogether deterring users. Having this in mind, we set to develop an easy, streamlined GUI that does not require any computing or image analysis skills, does not require installing additional software and that can rapidly analyze multiple images and organize the specific output. Our GUI provides an automated workflow that quantifies fibrosis of single or multiple histological images of skeletal muscle and heart tissue (Fig.1 and Supplementary Fig.1). To illustrate its functionality, we analyzed samples stained with SR, MT and IHC for collagen type I or collagen type III, all of which are commonly used to highlight fibrotic regions ^7,8^. The analyses are carried out in three distinct steps, each building upon the before ensuring comprehensive and accurate fibrosis quantification.

**Fig. I.**
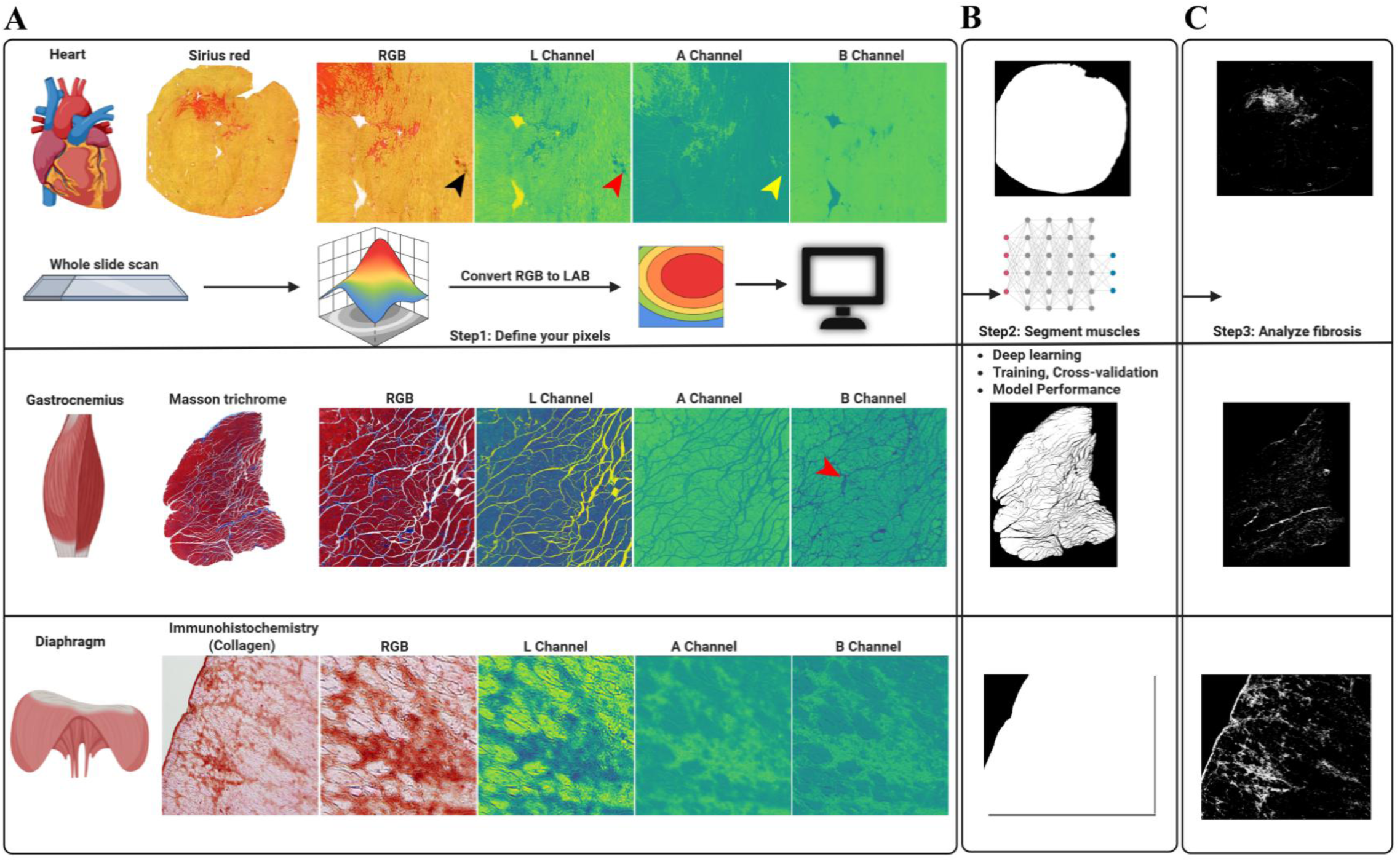
FibroTrack workflow for fibrosis analysis in skeletal and cardiac muscles. The workflow starts with whole-slide images stained with Sirius Red, Masson Trichrome, or Immunohistochemistry. **(A)** In Step **1,** Define Your Pixels, **RGB** images are converted to the **LAB** color space to facilitate user selection of pixel intensity values for identifying fibrotic regions. For Sirius Red staining, the A channel is primarily used for optimal fibrosis detection. While the L channel may show a similar pattern, it also highlights artifacts (red arrow in L channel, corresponding to black arrow in RGB), which the A channel effectively excludes (yellow arrow). For Masson Trichrome, fibrosis (dark blue-purple pixels indicated by red arrow) is most clearly visualized in the B channel, with limited fibrosis signal in the Land A channels. In Immunohistochemistry, fibrosis is visible across all LAB channels (L, A, and B), providing flexibility **for** the user to select either the A or B channel. **(B)** Step 2 Segment muscles, utilizes deep learning to segment the entire muscle tissue, distinguishing it from background. (C) In Step 3, Analyze Fibrosis, the software quantifies the fibrosis ratio percent within the segmented muscle.

First, when images are uploaded to FibroTrack, the software prompts users with a color normalization option (see color space conversion and normalization section). This critical step is highly recommended as it standardizes the color profiles across images, compensating for variations in staining intensity, microscope settings, and image acquisition parameters. After color normalization, FibroTrack automatically analyzes the images and suggests optimal pixel intensity threshold values for fibrotic tissue detection (see Step 1: Defining Pixel Values section). For SR-stained images, these values identify yellow-green hues representing fibrotic regions, while for MT-stained images, they isolate the blue-purple hues characteristic of fibrosis. Although the software provides these auto-selected values, users can manually adjust these thresholds if needed to accommodate unique sample characteristics or staining variations.

Next, FibroTrack employs a deep learning algorithm to perform whole muscle segmentation, a crucial step in distinguishing muscle tissue from the background. The model is trained on a diverse dataset of 2,034 images, which includes skeletal muscle and heart tissue images captured at various magnifications (e.g., 10x, 20x, 40x) and using different microscopes. This diversity ensures accurate segmentation, even across samples with varying staining intensities and protocols. The segmentation process also incorporates cross-validation to improve model performance and ensure consistent results.

Finally, fibrosis quantification is performed based on the segmented tissue areas. FibroTrack calculates the proportion of fibrotic tissue relative to the total muscle area, providing a precise and reliable measurement of fibrosis levels. This quantification is critical for assessing disease progression, comparing experimental conditions, and evaluating therapeutic interventions. The automation of this process ensures consistent and reproducible results, eliminating the variability associated with manual analysis. All generated data - including pixel intensity data, segmented images, and fibrosis ratios - are automatically saved to an output folder for easy access and further analysis.

### Color Space Conversion and normalization

Towards defining accurate pixel values, a color separation method is required. To do so, our GUI utilizes LAB color space. LAB is standardized by the International Commission on Illumination (CIE) ^9^ and ideal for color-based image analysis, as it separates lightness (L) from chromatic information (A and B channels). Channel A measures the green-to-red spectrum, while channel B represents the blue-to-yellow spectrum. This decoupling of lightness from chromaticity is particularly useful for analyzing histological images stained with SR or MT, where fibrotic tissue is typically highlighted in red or purple, respectively. For IHC images, fibrotic regions are similarly analyzed based on the module selected, red for SR or purple for MT (Fig.1A). Upon importing the folder containing the images, FibroTrack offers an automated color normalization feature (Supplementary Fig.2A) to address a common challenge in histopathology, the variability in staining intensities across tissue samples. This preprocessing step implements a robust statistical normalization algorithm that first calculates the RGB color distribution statistics for all images in the dataset. The software then identifies an optimal reference image whose color profile represents the median characteristics of the entire image set. Each image undergoes pixel-by-pixel transformation using a statistical formula that adjusts color values relative to this reference:

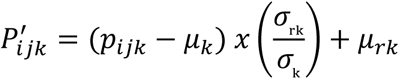

Where P’_ijk_ is the normalized pixel value at position (i,j) for color channel k, P_ijk_ is the original pixel value, μₖ and σₖ are the mean and standard deviation of channel k in the original image, and μᵣₖ and σᵣₖ are the mean and standard deviation of channel k in the reference image. This mathematical approach preserves the internal color relationships within each image while standardizing the overall color distribution across the dataset, ensuring that subsequent fibrosis detection operates on comparable color profiles regardless of original staining variations.

The choice of LAB color space over alternatives like HSV or RGB is critical for accurate fibrosis detection. Unlike HSV or RGB, LAB color space provides a more perceptually uniform representation that closely mimics human vision, allowing for better discrimination between fibrotic tissue and other tissue components or artifacts. In our comparative analysis, we found that the A channel (for SR staining) and B channel (for MT staining) in LAB space provide the most distinct separation between fibrotic tissue and other components (Supplementary Fig.2B,C), while HSV and RGB often detect artifacts or other tissue components (e.g., nuclei) as fibrosis, leading to inaccurate quantification (Supplementary Fig.3). For example, in SR-stained samples, the A channel effectively excludes artifacts (visible as dark regions in the L channel but not detected in the A channel), which would otherwise be misidentified as fibrosis in RGB-based analysis. Similarly, for MT staining, the B channel clearly visualizes fibrotic regions (dark blue-purple pixels) with minimal signal from non-fibrotic tissue, while other color spaces show insufficient contrast for reliable delineation. This superior specificity ensures that FibroTrack accurately identifies only fibrotic regions, even in images with variable staining quality or artifacts.

#### Step 1: Defining Pixel Values

Accurate fibrosis quantification in histological images requires precise identification of fibrotic regions, which are characterized by specific staining intensities. FibroTrack streamlines this process through an intelligent auto-selection algorithm that serves as the primary approach for determining optimal pixel thresholds. Upon image loading, the software automatically analyzes the LAB color space distribution according to staining type, with Sirius Red analysis focusing on brighter pixel values in the A channel (70th-99th percentiles) and Masson Trichrome targeting the darkest 5% of pixels in the B channel. Users are presented with these auto-selected values through a pop-up window (Supplementary Fig.4A, B), allowing immediate acceptance or optional manual refinement.

While the automatic selection is optimized for most standard preparations, the manual adjustment capability remains essential for handling variations in staining intensity that may occur due to differences in tissue preparation, staining protocols, and imaging conditions. Without this adjustment option, certain analyses could potentially misidentify tissue regions or fail to detect subtle fibrotic changes.

FibroTrack interface supports this process by allowing users to load images stained with SR, MT or IHC, supporting common formats such as “jpg,” “jpeg,” “png,” “tif,” and “tiff” (Supplementary Fig.5A, B). Once an image is loaded, the interface displays the source image alongside its LAB color space split, with channel A specifically used for pixel value definition. Channel A isolates the red-green chromatic range, which is particularly useful for distinguishing fibrotic tissue in SR-stained images (Supplementary Fig.5C).

To assist in pixel selection, the interface includes zoom in and out functionality, enabling users to magnify regions where manually hovering over the tissue may be challenging (Fig.2A). This feature is crucial for selecting specific areas of interest in highly detailed regions of the tissue, ensuring precise pixel value definition.

**Fig. 2.**
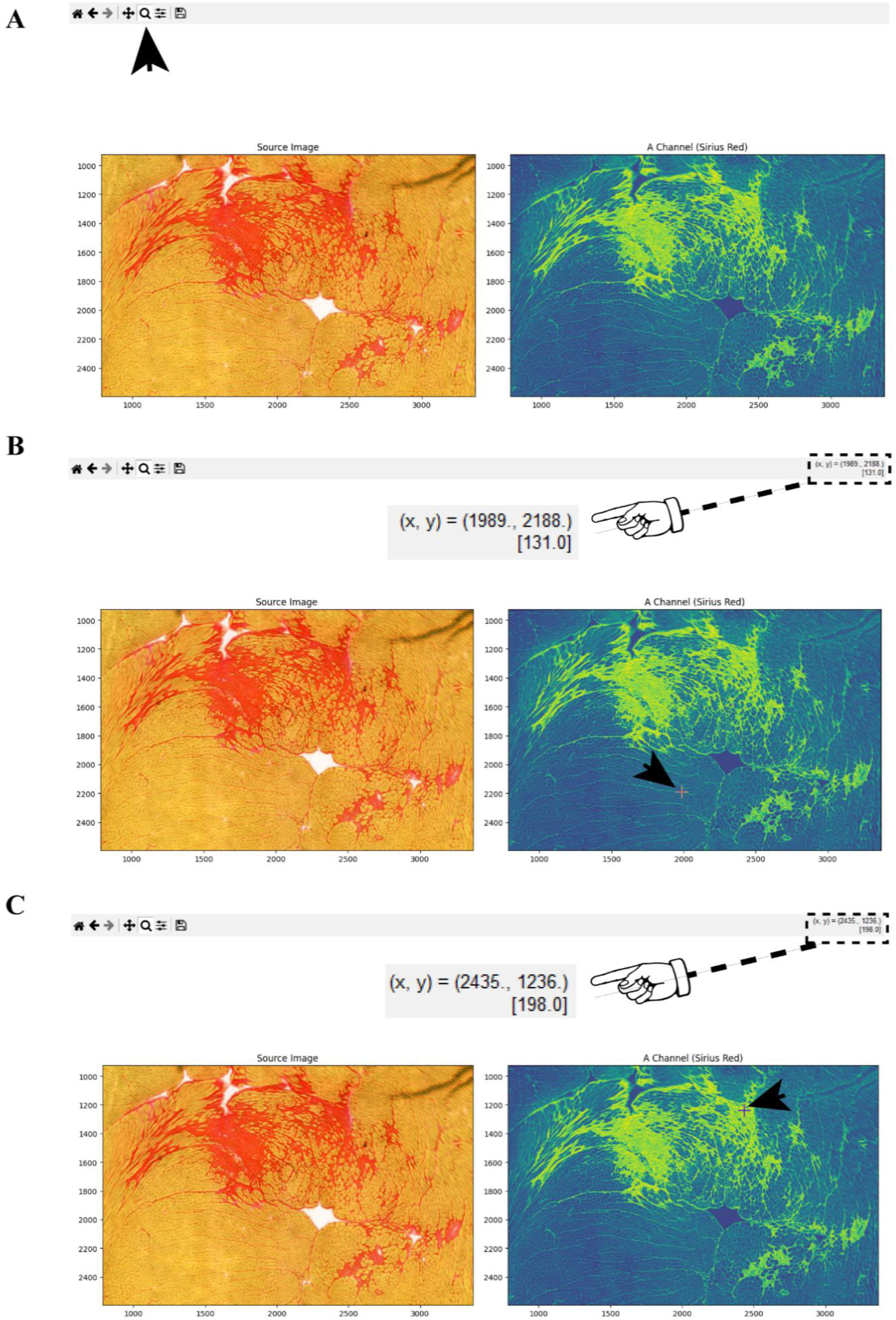
Defining lower and upper pixel intensity values for fibrotic and non-fibrotic tissue in Sirius Red-Stained Images. **(A)** The interface displaying the source image (left) and the corresponding LAB color space A channel (right). The zoom feature (indicated by the black arrow) allows users to magnify regions for accurate pixel selection. **(B)** The selection of the lower pixel intensity value [ 131.0], representing the darkest non-fibrotic area in the A channel. The black arrow points to the selected pixel location in the A channel, and the coordinates (x, y = 1989, 2188) of the selected pixel are shown in the interface. **(C)** The selection of the upper pixel intensity value [198.0], representing the brightest fibrotic area in the A channel. The black arrow highlights the selected pixel in the A channel, and the coordinates (x, y = 2435, 1236) of the selected pixel are displayed.

For the lower pixel value, the user picks the darkest yellowish hue within the A channel, which typically corresponds to the less fibrotic areas or the boundaries between fibrotic and non-fibrotic tissue. The pixel intensity value is displayed in the top right corner of the interface alongside the (x, y) coordinates of the cursor (Fig. 2B). Here, in the chosen sample, the lower pixel value is [131.0]. The user enters this value into the designated field labeled as Lower Pixel Value (Supplementary Fig.5D).

Next, the upper pixel value is defined by selecting the brightest yellowish region within the A channel, representing the most fibrotic areas. The pixel intensity value is recorded, and in this example, the upper value is [198.0] (Fig.2C). This value is entered into the Upper Pixel Value field (Supplementary Fig.5D).

After defining both the lower- and upper-pixel values, the user clicks the Save Pixel Values button (Supplementary Fig.5D). This action exports the pixel values as an Excel file named “Defined_pixel_values.xlsx,” saved in the same directory of the original selected image. The exported file contains the image name, the lower- and upper-pixel values, and the type of staining used (Supplementary Fig.5E). This feature ensures that users can easily track the pixel values applied on the selected image, enabling consistency across multiple samples and analyses. The same process can be applied to the MT and IHC-stained images (Supplementary Fig.6 and Supplementary Fig.7 respectively). To enhance user experience and reduce manual steps, we have implemented an automatic pixel value transfer feature to FibroTrack. This improvement allows pixel values selected in Step 1 to be automatically transferred to Step 3, eliminating the need for users to manually record and re-enter these values. This automation not only streamlines the workflow but further improves the reliability and efficiency of the analysis process.

#### Step 2: Muscle Segmentation and model performance

Proper segmentation of muscle tissue is among the most important prerequisites for quantitative fibrosis ratios^10–12^. The fibrosis ratio (*F_R_*) is mathematically defined as the proportion of fibrotic tissue area (*A_f_*) relative to the total muscle area (*A_m_*) and can be expressed as:

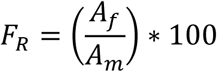

The use of a deep learning model for tissue segmentation provides significant advantages over simpler methods like binarization or thresholding. While binarization might seem sufficient for distinguishing tissue from background, it is highly sensitive to variations in staining intensity and background noise, often leading to inaccurate results (Supplementary Fig.8A).

Further, ImageJ is widely used for biological image analysis, however it presents significant limitations for fibrosis quantification. As demonstrated in Supplementary Fig.8A, ImageJ’s automatic thresholding struggles and presents critical disadvantages in distinguishing true fibrosis from artifacts in heart tissue samples, where staining inconsistencies can create artifacts that closely resemble fibrotic tissue under standard parameters.

Similarly, Supplementary Fig.8B illustrates how commonly recommended ImageJ color deconvolution protocols may deliver less accurate and less efficient results compared to tailored thresholding techniques for cardiac fibrosis quantification. When tissue samples were processed through standard color deconvolution vectors (RGB vectors for Sirius Red staining and Masson’s Trichrome vectors for Masson’s Trichrome staining), fibrosis detection was substantially inferior compared to our custom thresholding approach.

Furthermore, implementation barriers can significantly impact research efficiency. As shown in Supplementary Fig.9, alternative solutions such as FIBER-ML face substantial technical hurdles, including specific MATLAB version dependencies (v2019b or earlier), plus Image Toolbox and Statistical ML Toolbox installations. These requirements create significant obstacles to accessibility, with users encountering multiple execution failures (evidenced by the error messages shown). By contrast, FibroTrack offers a standalone solution requiring no programming knowledge and no software dependencies, making it immediately accessible to researchers across varying technical backgrounds.

To address these limitations, FibroTrack employs a deep learning-based model built on the YOLOv11 (You Only Look Once, version 11) framework^13^, a highly efficient and state-of-the-art architecture designed for both object detection and image segmentation tasks. YOLOv11 architecture is known for its balance between speed and accuracy, making it particularly suitable for handling large datasets while maintaining precise segmentation (Fig. 3).

**Fig. 3.**
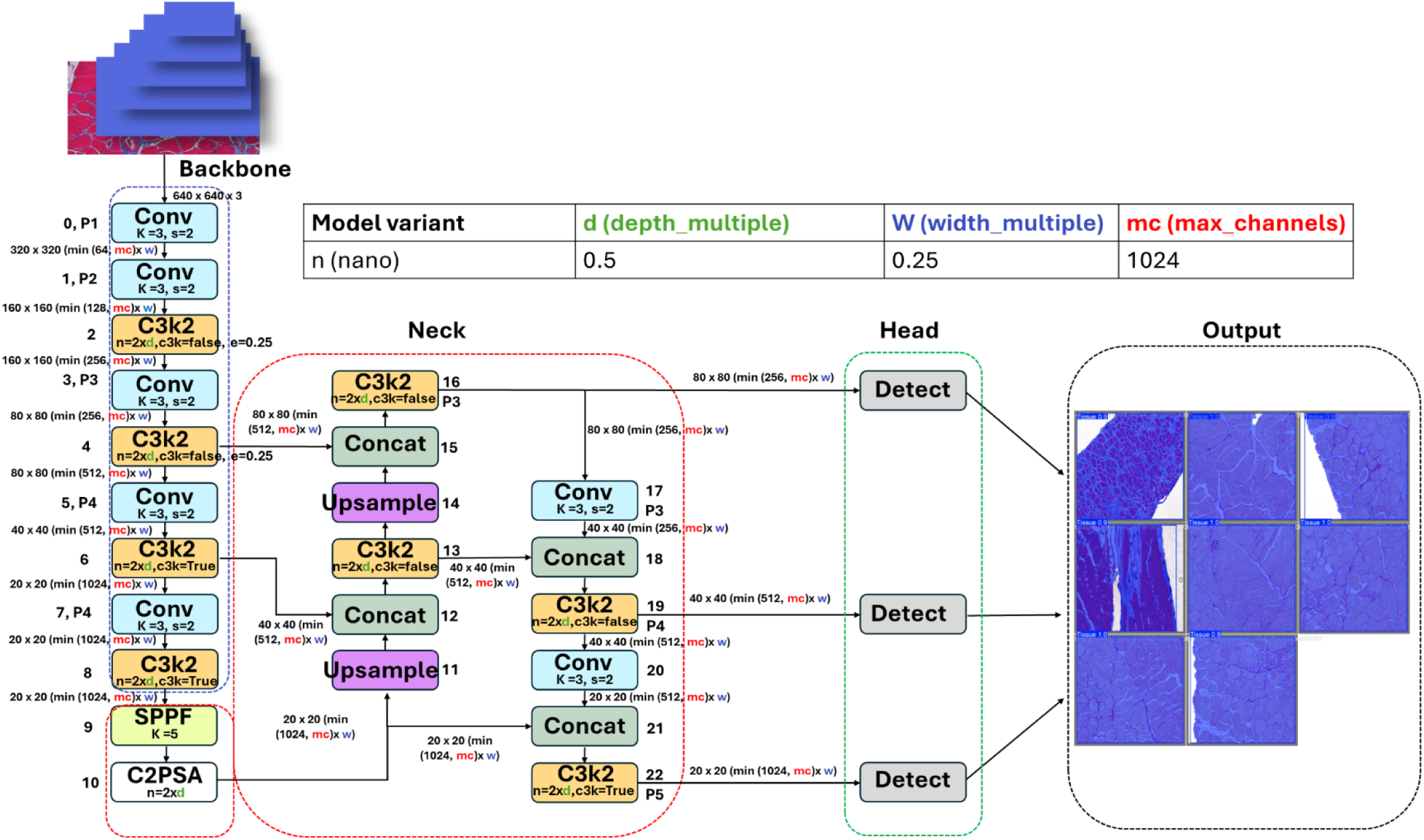
FibroTrack’s Mode] YOLOVll Architecture. **(A)** Input image and the backbone network based on convolutional layers and C3K.2 modules, extracting features at different scales. **(B)** The Neck section combines features through concatenation and upsampling operations. **(C)** The Head performs detection at three different scales, generating output segmentations. **(D)** The table defines the parameters for the “nano” model variant used in YOLOvl 1, including depth and width multipliers, and maximum channels. **(E)** Output showing segmented muscle tissue images.

YOLOv11, trained on diverse tissue samples, is more robust to these variations and can accurately delineate complex tissue boundaries, even in the presence of artifacts or irregular tissue shapes, addressing the limitations highlighted in the comparative analyses.

The training dataset used for developing the YOLOv11 model consisted of 2034 histological images stained with SR and MT. These images included cardiac and skeletal muscles with distinct staining intensities, and experimental conditions, ensuring the model’s ability to generalize across different datasets. Each image was pre-processed with auto-orientation of pixel data (with EXIF-orientation stripping) and resized to 640×640 (stretch). To further enhance the model’s robustness, data augmentation techniques were applied for each source image, including random brightness adjustment (±25%), random exposure adjustment (±14%), and random Gaussian blur of between 0 and 2.5 pixels. This process expanded the dataset to 4650 images, significantly improving the model’s ability to handle variability in staining protocols and imaging conditions. The dataset was annotated and augmented using Roboflow^14^, a widely used platform for optimizing machine learning datasets. The dataset used for training the model was split into three parts: 70% (training), 20% (validation), and 10% (testing), ensuring a well-rounded evaluation of the model’s generalization ability (Fig.4). Concerning this aim, key metrics parameters were derived from the YOLOv11 validation dataset. The *box precision (Box (P))* metrics refers to the percentage of predicted muscle regions (bounding box) that are correct, reached a precision of 95.6% (Table 1 and Supplementary Table 1).

**Fig. 4.**
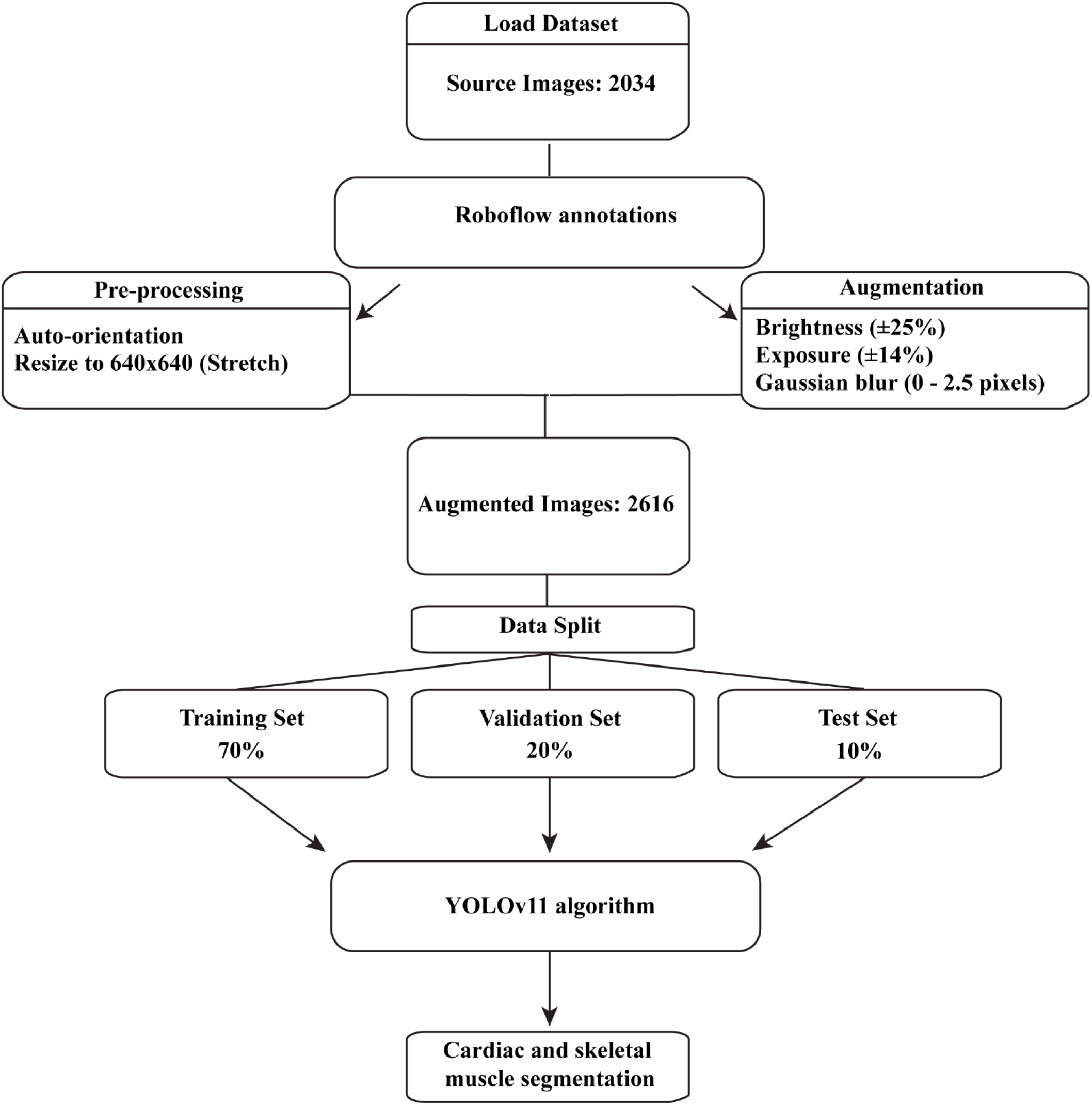
Flowchart illustrating the steps for dataset preparation and processing. The process commence with loading and annotating the dataset, followed by pre-processing and augmentation. The dataset is then split into training, validation, and test sets, before applying the YOLOvl 1 algorithm for cardiac and skeletal muscle segmentation.

**Table 1.** YOLOvll Model Validation Summary. Comprehensive summary of the YOLOvl 1 n-seg model validation metrics. The Model Information section includes details about the framework (Ultralytics YOLOvll), hardware used (Quadro RTX 5000, 16,384 MiB), and environment specifications (Python 3.12.3, PyTorch 2.5.0, CUDA device 0). The Performance section reports the precision (P), recall (R), and mean average precision (mAP50-95) for both bounding boxes and segmentation masks. Bounding box precision was (95.6%), recall was (97.8%), and (mAP 50-95) was (97.4%). For segmentation masks, the precision was (97.5%), recall was (95.3%), and (mAP50-95) was (97.2%). The Speed section highlights the model’s computational efficiency, with preprocessing, inference, and postprocessing times averaging 0.3 ms, 3.7 ms, and 1.2 ms, respectively. The Training Details show that the model completed 114 epochs in a total training time of 10 minutes, with a processing speed of 11.39 images per second. Finally, the Results Location indicates the saved model weights best.pt as well as last.pt, representing the best performing checkpoint during training.

| Category | Metric | Value |
| --- | --- | --- |
| Modle Information | Model | YOLOv11n-seg |
|  | Framework | Ultralytics YOLOv11 |
|  | Hardware | Quadro RTX 5000 (16,384 MiB) |
|  | Python Version | 3.12.3 |
|  | PyTorch Version | 2.5.0 |
|  | CUDA Device | 0 |
| Performance | Box Precision (P) | 0.956 (95.6%) |
|  | Box Recall (R) | 0.978 (97.8%) |
|  | Box mAP<sub>(50-90)</sub> | 0.974 (97.4%) |
|  | Mask Precision (P) | 0.975 (97.5%) |
|  | Mask Recall (R) | 0.953 (95.3%) |
|  | Mask mAP<sub>(50-90)</sub> | 0.972 (97.2%) |
| Speed | Preprocessing Time | 0.3 ms |
|  | Inference Time | 3.7 ms |
|  | Loss Calculation Time | 0.0 ms |
|  | Postprocessing Time | 1.2 ms |
| Training Details | Epochs Completed | 114/114 |
|  | Total Training Time | 00:10:00 |
|  | Processing Speed | 11.39 images/sec |
| Results Location | Saved Weights | best.pt |
|  |  | last.pt |

A precision of 95.6% indicates that nearly all the regions the model classified as muscle tissue were indeed correct. Similarly, the *Box Recall (Box(R))* measures the percentage of the actual regions (ground truth) that were successfully detected by the model. This metric resulted in a higher recall value of (0.978), Suggesting that 97.8% of the actual tissue regions present in the images were correctly detected by the model’s bounding boxes, minimizing false negatives (Table 1 and Supplementary Table 1).

The recall (R) and precision (P) are calculated according to the following equations:

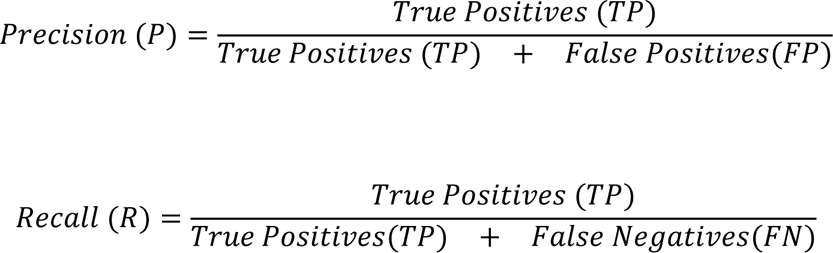

The *Mean Average Precision (mAP50)* at *Intersection over Union (IoU)* 0.5 is a standard metric used to evaluate object detection models. It calculates the average precision across all classes at a specific threshold for IoU. This IoU is a measure of how much the area of predicted bounding box overlaps with the area of the ground truth box. In this case, mAP50 of 0.974 means that the model achieved 97.4% average precision when the overlap between the predicted and actual bounding boxes is at least 50% (IoU ≥ 0.5) (Table 1 and Supplementary Table 1). The Intersection over Union (IoU) Formula is defined by the following equation:

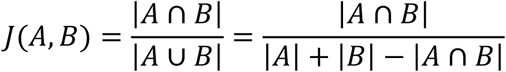

Where A represents the predicted bounding box, and B represents the ground truth box. The higher the value of the IoU, the higher the model performance is.

Another important metric is *mAP50-95*. This is a more comprehensive version of mAP50 that averages the precision over a range of IoU thresholds, from 0.5 to 0.95 in increments of 0.05. This metric is stricter since it requires good performance across varying levels of overlap between the predicted and actual bounding boxes. mAP50-95 of 0.972 means that the model achieved an average precision of 97.2% across IoU thresholds from 50% to 95% (Table 1 and Supplementary Table 1). This indicates that the model performs consistently well even when the overlap between the predicted and actual regions is required to be very high.

In terms of pixel-level segmentation, the model performed equally well, with a *Mask Precision* of 97.5% and a *Mask Recall* of 95.3%, demonstrating its high accuracy in correctly identifying and segmenting muscle tissue pixels (Table 1 and Supplementary Table 1). Finally, the model’s efficiency is also notable, processing images in just 0.3ms for *preprocessing*, 3.7 milliseconds (ms) for *inference*, and 1.2ms for *postprocessing*, totaling roughly 5.2 ms per image, making it highly suitable for handling large datasets. The model was trained in Python, with the best-performing version saved as ‘best.pt’, based on validation accuracy and segmentation performance (Table 1).

To evaluate the consistency of our model’s performance across different samples, we analyzed the error distribution on our test dataset (n=406). The confusion matrix demonstrates high segmentation performance with 96% of tissue pixels correctly identified (sensitivity/true positive rate), while achieving perfect 100% specificity for background (true negative rate). Only 4% of tissue pixels were incorrectly classified as background (false negative rate), with no background pixels misclassified as tissue (0% false positive rate) (Supplementary Fig.10A). The model achieved F1-score of 97%, indicating highly accurate overlap between predicted and ground truth segmentation masks. Analysis of the spatial distribution of model predictions confirms that segmentation accuracy is maintained regardless of the size, shape, or position of tissue within the images, with consistent performance across both positional coordinates and dimensional variations (Supplementary Fig.10B). This spatial consistency across diverse tissue morphologies indicates that FibroTrack’s segmentation capabilities are robust and not disproportionately affected by specific tissue configurations, ensuring reliable quantification results across different sample types and staining conditions.

These metrics collectively show that the YOLOv11-based segmentation in FibroTrack is both - highly accurate and efficient, enabling precise and reliable fibrosis quantification with minimal user intervention (Supplementary Figs.11, 12 and Supplementary Table 1).

FibroTrack’s user interface simplifies the segmentation process, allowing users to easily load the pre-trained YOLOv11 model and select a folder containing the images to be segmented (Fig.5A). Users begin by clicking the “Load YOLO Model” button, which prompts them to select the ‘best.pt’ file (Fig.5B). After loading the model, users can choose the folder containing the SR or MT-stained images for analysis (Fig.5C).

**Fig. 5.**
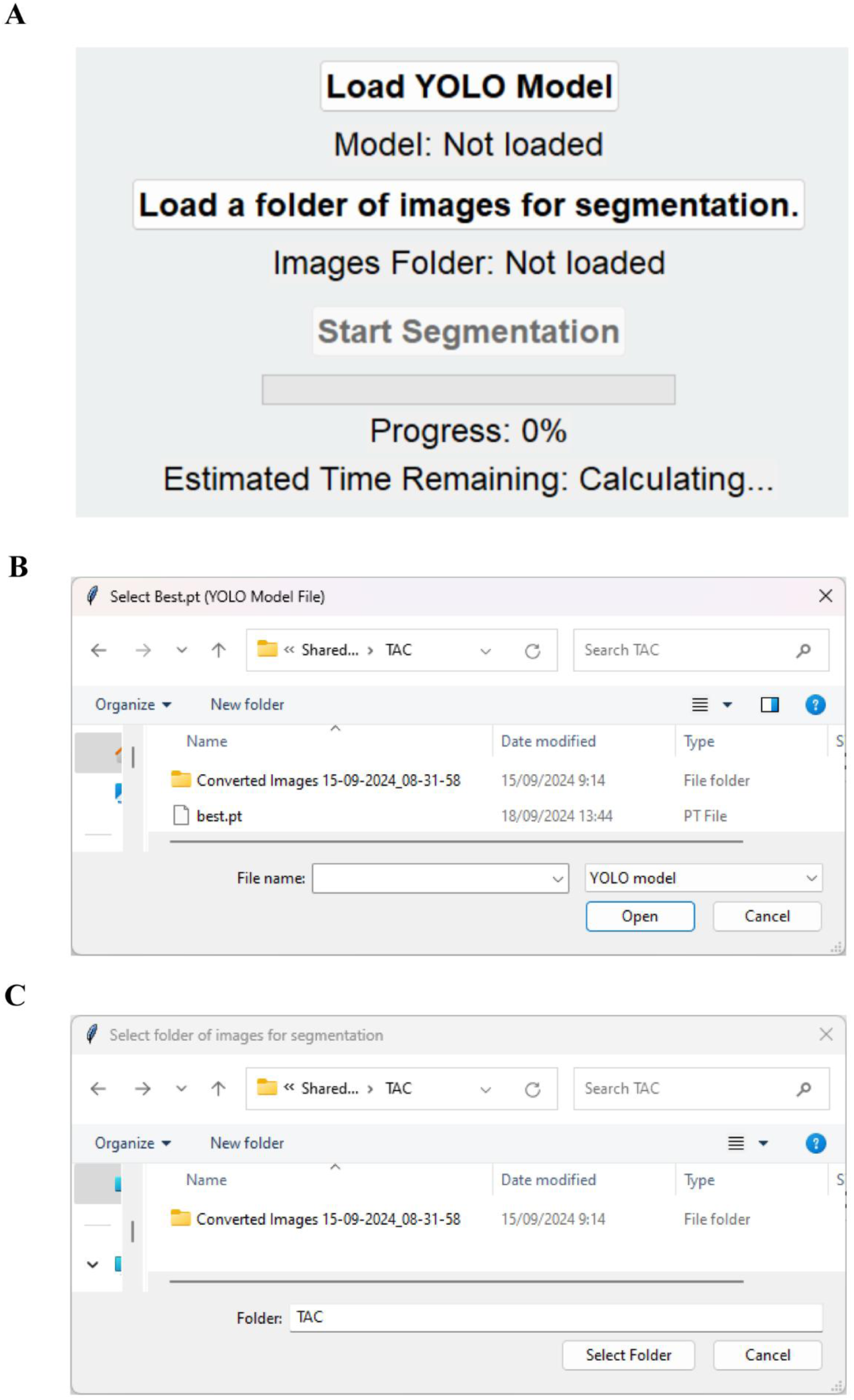
FibroTrack User Interface for Segmentation Workflow. **(A)** Users can load the pre-trained YOLOvl **1** model and select a folder containing images for segmentation. By doing so, the Start Segmentation button becomes active, enabling users to initiate the segmentation process. The interface clearly displays the status of the loaded model and selected image folder, along with a progress bar and an estimated time remaining for the segmentation process. **(B)** File selection dialog prompting the user to load the YOLOv8 model weights (best.pt file) from a directory. This file contains the pre-trained model required for the segmentation task. **(C)** File selection dialog where the user selects a folder containing the Sirius Red or Masson Trichrome-stained images to be segmented. Once selected, the images in this folder are auto­ matically processed by FibroTrack.

Once the segmentation process is initiated, by clicking the “Start Segmentation” button, FibroTrack automatically processes all images in the selected directory. The tool supports a wide range of image formats, including JPG, PNG, and TIFF, ensuring compatibility with commonly used file types in histological analysis. While users do not see individual images during the folder selection process, all images within the chosen directory are imported and segmented automatically.

After segmentation is completed, segmented images are stored in four main subfolders, each serving a specific purpose: i) “Binary_masks_YOLOV11_results” contains binary mask representations of the segmented tissue regions and an Excel file named “Muscle_Region_Properties_YOLOV11.xlsx” with detailed quantitative properties; ii) “Removed_blood_vessels_and_inner_empty_chamber” houses processed images where artifacts such as empty heart chambers and interstitial spaces between muscle fibers (resulting from fixation or technical processing) have been removed to prevent skewing fibrosis quantification results, including an Excel file with adjusted measurements and a subfolder containing the corresponding binary masks; iii) “Segmented_images_with_bbox” includes segmented images with bounding boxes overlaid for easy visualization of detected regions; and iv) “Segmented_images_without_bbox” stores segmented images without bounding boxes for a cleaner representation. This enhanced output structure ensures users can access both raw segmentation results and refined data where technical artifacts have been addressed, providing a more accurate foundation for downstream quantitative analysis of tissue fibrosis. In addition to its powerful segmentation capabilities, FibroTrack incorporates a real-time progress bar and an estimated time remaining feature, providing users with continuous feedback throughout the segmentation process. The progress bar visually indicates the status of the segmentation task, while the estimated time remaining helps users manage their workflow efficiently, particularly in large-scale studies where processing times can be substantial. This feature is especially useful for researchers handling large datasets, as it allows for better time management and resource allocation.

By leveraging the YOLOv11 model, FibroTrack ensures accurate and efficient segmentation of muscle tissue, significantly reducing the variability associated with manual annotation and enhancing the reproducibility of fibrosis quantification across different samples. The combination of an advanced deep learning model and a user-friendly interface makes FibroTrack a highly effective tool for large-scale fibrosis studies, providing consistent results with minimal user input.

#### Step 3: Analyze Fibrosis

In this final stage, the segmented images from the previous step are ready for fibrosis ratio quantification. Users begin by uploading the segmented images from the “segmented_images_without_bbox” folder. The interface allows for quick and easy upload, streamlining the start of the analysis process (Fig.6A).

**Fig. 6.**
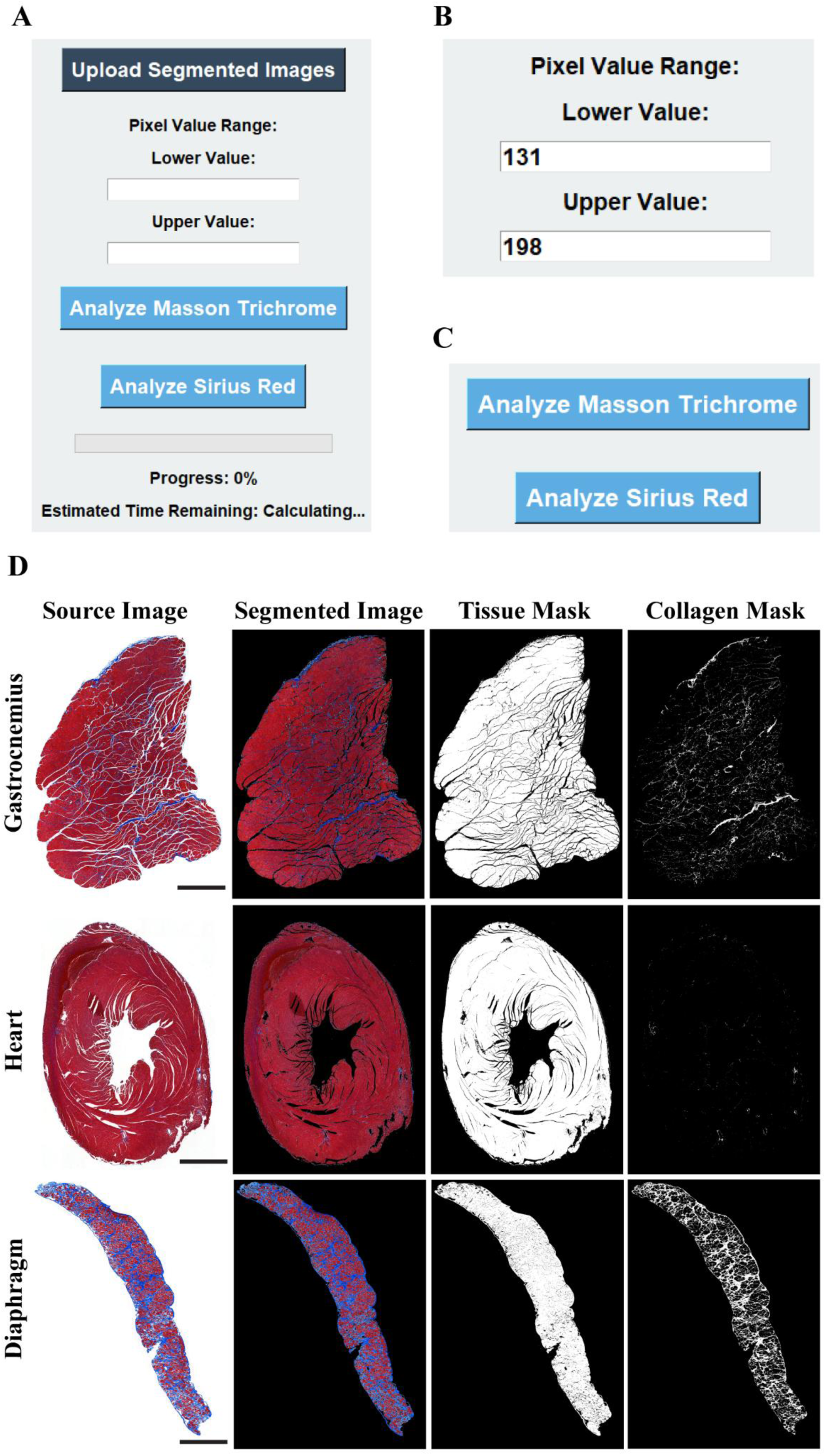
FibroTrack Interface and Analysis Workflow: Comprehensive Fibrosis Mapping in Cardiac and Skeletal Muscles. **(A)** Main interface for uploading segmented images and defining the pixel value range for fibrosis detection. Users can initiate the analysis by selecting either Sirius Red or Masson Trichrome staining. **(B)** Input fields for specifying the lower- and upper-pixel value thresholds to accurately identify fibrotic tissue. **(C)** Buttons to select the approprate analysis method based on the stain used, Sirius Red or Masson Trichrome. **(D)** Representive images of analyzed cardiac and skeletal muscles. Scale bar: 200µm.

Once the images are uploaded, users are required to input the pixel value range. This range, consisting of upper and lower limits, is crucial to defining the fibrosis based on the specific staining used. The values should be referenced from the “Defined_pixel_values.xlsx” file generated earlier; Step 1: Defining Pixel Values (Supplementary Fig.5D) to maintain consistency. Accurately setting these pixel limits ensures that the analysis correctly distinguishes between fibrotic tissue and muscle tissue, avoiding any misclassification that could skew the results.

After setting the pixel values (Fig.6B), users choose between “Analyze Sirius Red” or “Analyze Masson Trichrome” based on the stain used in their experiment (Fig.6C). Similarly, for IHC staining, users can choose either the SR or the MT modules to analyze the fibrotic regions. This selection initiates the analysis of where the GUI calculates the fibrosis ratio by identifying the collagen area and normalizing it relative to the total muscle area (Fig.6D). This process runs automatically once initiated.

As the analysis proceeds, a progress bar updates in real-time, providing users with a visual indication of how far along the analysis is.

Upon completion, the analysis results are saved in a new subfolder within the “segmented_images_without_bbox “ directory, named Final_Results_Analyzed_Fibrotic_Muscles_{day_name}_{date_time}, ensuring easy identification and organization of the output. This timestamped subfolder contains the segmented images and a detailed CSV file, named Fibrosis (%) Results{channel}{day_name}{date_time}.csv. The CSV file includes crucial data such as the filename of each image, the collagen area, the total muscle area, and the fibrosis ratio expressed as a percentage, allowing for straightforward analysis and reference. The CSV file follows a naming convention that reflects the type of analysis performed and when it was run. It is structured to allow researchers to quickly navigate through the data, with key metrics for each image clearly outlined. This organized format enables efficient tracking of fibrosis across all images and ensures that critical information is preserved for further analysis.

By automating the creation of both, the results folder and the corresponding CSV file, the tool eliminates the need for manual data sorting or entry, significantly reducing the chance for errors. This also enhances the reproducibility and traceability of the results, as all data is saved in a consistent, easily accessible manner.

The entire process - from uploading segmented images to generating the final fibrosis metrics - is designed to be user-friendly while maintaining accuracy. The clear interface, real-time progress updates, and structured output ensure a smooth workflow for researchers, even when dealing with large-scale studies.

### Validation Against Independent Pathologists’ Assessments

To validate FibroTrack’s accuracy, we compared its automated fibrosis quantification with manual assessments by two independent pathologists (P1 and P2), each performed on the same set of 31 muscle tissue samples exhibiting varying degrees of fibrosis. Importantly, all evaluations were performed in a blind manner, with pathologists unaware of FibroTrack’s measurements and each other’s assessments.

The results demonstrate strong consistency between FibroTrack’s fibrosis quantification (mean 18.4%, median 21.4%) and the assessments by P1 (mean 16.8%, median 20.0%) and P2 (mean 17.6%, median 20.0%) (Fig. 7A). Statistical analysis using the Kruskal-Wallis test revealed no significant difference between the three measurement methods (p = 0.96), confirming that our automated approach produces results consistent with pathologists’ evaluations.

**Fig. 7.**
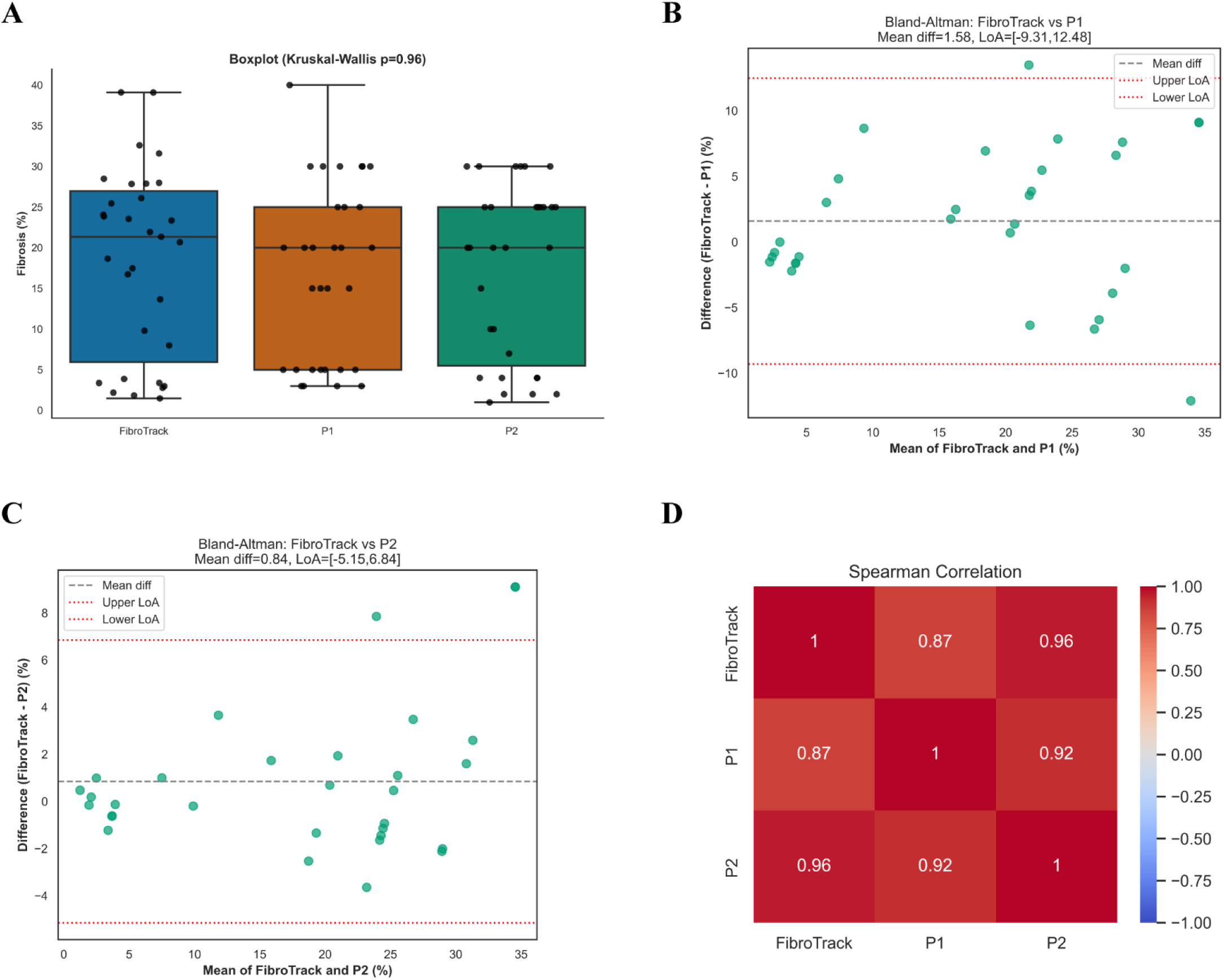
Validation of FibroTrack Against Independent Pathologists’ Assessments. **(A)** Boxplot comparing fibrosis quantification(%) across the three assessment methods: FibroTrack, first pathologist (PI), and second pathologist (P2). The Kruskal-Wallis test (p=0.96) shows no significant difference between the three methods. Each dot represents an individual sample measurement, with median values indicated by horizontal lines within boxes. **(B)** Bland-Altman plot comparing FibroTrack with the first pathologist (Pl). The plot shows the difference between FibroTrack and P1 measurements (y-axis) against their mean (x-axis). The mean difference (1.58%, dashed grey line) and limits of agreement (-9.31% to 12.48%, dotted red lines) demonstrate strong clinical consistency . **(C)** Bland-Altman plot comparing FibroTrack with the second pathologist (P2). The mean difference (0.84%, dashed grey line) and narrower limits of agreement (-5.15% to 6.84%, dotted red lines) indicate excellent agreement between FibroTrack and P2 assessments. **(D)** Heatmap of Spearman correlation coefficients between the three assessment methods. The strong correlations between FibroTrack and both pathologists (r=0.87 with Pl, r=0.96 with P2) demonstrate that FibroTrack not only produces similar measurements but also ranks samples consistently with expert pathologists.

Further analysis using Bland-Altman plots shows also agreement between FibroTrack and both pathologists’ assessments. The mean difference between FibroTrack and P1 was 1.58% (limits of agreement: -9.31% to 12.48%) (Fig. 7B), while the mean difference with P2 was even smaller at 0.84% (limits of agreement: -5.15% to 6.84%) (Fig. 7C). These narrow limits of agreement indicate strong clinical consistency between the automated and manual assessments.

Most importantly, Spearman correlation analysis revealed a strong correlation between FibroTrack and both pathologists (r = 0.87 with P1 and r = 0.96 with P2) (Fig. 7D). This high degree of correlation demonstrates that FibroTrack not only produces similar overall fibrosis measurements but also ranks samples consistently with two independent pathologists across the spectrum of fibrosis severity.

Notably, while the median values were similar across all three assessment methods, FibroTrack showed a slightly wider distribution of values (range: 1.5% - 39.1%) compared to both pathologists (P1 range: 3% - 40% , and P2 range: 1% - 30%). This suggests that FibroTrack may detect subtle variations in fibrosis that might be normalized during human assessments. The consistency of central tendency metrics across all three methods, despite this slight difference in range, validates our approach as a reliable alternative to manual expert assessments.

This validation is particularly important as it demonstrates that FibroTrack can achieve high-level accuracy while eliminating the subjectivity and time constraints inherent to manual analysis. Furthermore, the consistency between automated and pathologists’ assessments supports FibroTrack’s utility as a standardized tool for fibrosis quantification across different research environments.

## Discussion

We developed FibroTrack, a fully automated platform for fibrosis quantification in skeletal and cardiac muscle histology. The tool combines LAB color space normalization with the YOLOv11 deep learning architecture, addressing two persistent challenges in fibrosis analysis: (i) variability in staining and (ii) the need for reliable segmentation of complex tissue morphologies. This integration is the central novelty of our work. Previous methods have typically relied either on color analysis or on segmentation algorithms, often requiring programming expertise, external toolboxes, or manual adjustment, which limits reproducibility and accessibility^4^. By contrast, FibroTrack unites both strategies in a standalone graphical interface, making advanced fibrosis quantification available to a wider range of researchers.

The choice of LAB color space was crucial. Compared with HSV and RGB, LAB provided superior separation of fibrotic tissue from background and staining artifacts (Supplementary Figs. 2,3 and 8). In direct comparisons, ImageJ’s color deconvolution methods frequently failed to distinguish fibrosis reliably in Sirius Red and Masson’s Trichrome stained samples, especially when staining intensity varied between experiments. FibroTrack maintained accurate detection under these conditions, highlighting the robustness of LAB-based approaches. We also compared our system with FIBER-ML^4^, a machine-learning tool that requires MATLAB and additional packages. While effective, its reliance on specialized software and coding knowledge limits adoption. FibroTrack avoids these barriers by operating as a standalone application that requires no programming skills or external dependencies, offering a clear usability advantage.

In addition to color normalization, FibroTrack incorporates a deep learning segmentation module powered by YOLOv11. Our model was trained on 2,034 annotated histological images spanning diverse tissue types, staining intensities, and experimental conditions. To enhance robustness, training included targeted data augmentation such as random changes in exposure, brightness, and image blurriness, simulating the variability commonly encountered in histological preparations and image acquisition (Fig.4). This design ensures consistent accuracy across protocols and laboratories, making the system resilient to real-world challenges of tissue preparation and imaging.

Validation against blinded pathologists confirmed the accuracy of FibroTrack. Correlations with manual scoring reached r = 0.96, and no statistically significant differences were observed between automated and expert assessments. Interestingly, FibroTrack captured subtle variations in fibrosis that were often normalized during manual evaluation, suggesting that computational analysis may provide greater sensitivity while eliminating inter-observer variability. These findings position FibroTrack not as a replacement for expert pathologists, but as a complementary tool that improves reproducibility and supports standardized analysis.

The biological and clinical relevance of this work is considerable. Fibrosis is a defining feature of Duchenne muscular dystrophy and other muscular dystrophies^15^, where up to 40-50% of muscle tissue may be replaced by fibrotic tissue in advanced stages^16^. Similarly, cardiac fibrosis contributes to pathological remodelling, arrhythmias, and progression to heart failure, and is strongly associated with worse clinical outcomes^17^. Despite its importance, fibrosis quantification remains inconsistent and time-consuming, particularly in studies requiring high throughput. Manual methods are laborious and prone to observer bias, while existing automated tools often depend on technical expertise or produce variable results across staining protocols. FibroTrack directly addresses these limitations by providing an accurate, reproducible, and scalable solution that reduces workload while maintaining clinical-grade performance.

The software is also designed with usability in mind. Automated outputs include segmented images and structured spreadsheets of fibrosis metrics, organized into timestamped folders for straightforward traceability. Processing speed of approximately 5 milliseconds per image makes FibroTrack suitable for large-scale studies, while real-time progress feedback further streamlines workflow management. These features make FibroTrack practical for both small laboratories and multicentred collaborations, ensuring that data can be generated rapidly and consistently across diverse settings.

In summary, FibroTrack represents a significant advance in fibrosis quantification by uniquely combining LAB color normalization with YOLOv11 segmentation in a user-friendly standalone platform. The system achieves high accuracy and reproducibility, eliminates observer bias, and scales efficiently for large studies. By minimizing technical barriers and standardizing fibrosis analysis, FibroTrack offers a novel and clinically relevant tool that has the potential to accelerate preclinical research, support translational studies, and contribute to more consistent pathology workflows.

### Declaration of generative AI and AI-assisted technologies in the writing process

During the preparation of this work the authors used Claude 3.5 Sonnet to improve the readability of the manuscript. After using this Claude 3.5 Sonnet, the authors reviewed and edited the content as needed and took full responsibility for the content of the published article.

## Methods

### Automatic slide scanner

SR and MT Slides were digitalized using a Panoramic Flash 250 digital slide scanner (3DHistech, Budapest, Hungary) in bright-field mode using a x20/0.8 objective lens, pixel size 0.242nm with the extended focus mode to select the sharpest image from 3 focal planes spaced 1 μm apart.

### Olympus VS200 digital scanner - Brightfield

SR and MT Tissue sections were digitally scanned with the Olympus VS200 digital scanner (Olympus Corp., Tokyo, Japan) with a 20x objective lens at a resolution of 0.2730 μm/pixel and saved as vsi files.

### OLYMPUS cellSens Standard

SR and MT slides were captured at various magnifications with objectives of 10x, 20x, 40x at a resolution of 0.645 μm/pixel, 0.3225 μm/pixel and 0.16125 μm/pixel respectively.

### Hardware and GUI Configuration for Model Training

The deep learning model, YOLOv11n-seg, was trained on a high-performance machine (HIVE3049) with an Intel(R) Xeon(R) Silver 4210R CPU @ 2.40GHz (40 processors), 512 GB RAM, and a Quadro RTX 5000 GPU with 16,384 MiB memory. The system ran a 64-bit, x64-based operating system. The training was performed using Ultralytics YOLOv11. Framework, implemented in Python 3.12.4 and PyTorch 2.0.4, with GPU acceleration enabled via CUDA (device 0). The training process utilizes 2,000 epochs with a batch size of 4 and an image size of 640 pixels. During training, the dataset was loaded into RAM to enhance performance, and the optimizer was set to automatic configuration. Additionally, an initial learning rate of (1×10^−4^), a final learning rate of (1×10^−3^) and a momentum of (9.37×10^−1^) were used, with weight decay configured at (5×10^−4^). Data augmentation techniques such as random translation (0.1), scaling (0.5), horizontal flipping (0.5), and mosaic augmentation (1.0) were applied to improve model generalization. The model was evaluated using an Intersection over Union (IoU) threshold of 0.7 and overlap masks with a ratio of 4 were employed. Training results were saved to the project directory, and simplifications in the model were applied using the torchscript format. The training process was executed deterministically, ensuring reproducibility across runs.

### Recommended System Requirements for FibroTrack

For optimal performance of FibroTrack, the following system specifications are recommended: Operating System: Windows 10/11 Pro 64-bit or Windows Server 2019 Standard. Processor: 11th Gen Intel Core i7 or better (2.50GHz or higher). RAM: 16 GB ,Storage: 658.2 MB available space.

### Data availability

The FibroTrack raw image data, dataset splits into training, validation and test sets, annotation details, and YOLOv11 configuration files optimized for building a segmentation model are all provided at https://github.com/Anas-Odeh/FibroTrack. Specifically, the raw images are in FibroTrack_Raw_Images_yolov11.rar, while train.rar, valid.rar and test.rar contain the separated training, validation and test splits respectively.

### Code availability

The custom analysis GUI is described at https://github.com/Anas-Odeh/FibroTrack, and the source python code by the name of “FibroTrackSourceCodeFinal.py” under MIT license.

### Software Installation

The FibroTrack graphical user interface (GUI) installation package is accessible through Google Drive(https://drive.google.com/file/d/1PvTxR_7k43wXrBx1fXlQomSxXSApQrA2/view?usp=sharing). The installation process requires downloading the software package, followed by extracting the “ FibroTrack installation.zip” folder from the downloaded package. The installation is completed by executing the “Install FibroTrack Graphical User Interface.exe” file, which sets up the complete GUI system. This installation procedure ensures proper configuration of all necessary components for utilizing the FibroTrack system, including the user interface elements and associated icons. (See GitHub, Guidelines for FibroTrack Installation software For Windows Users).

## Acknowledgements

We would like to thank Lama Awwad, Laris Achlaug, Anna Kaganovsky and Ami Aronheim for providing histological slides stained with Sirius red and masson trichrome. We would like to thank Maya Holdengreber and Melia Gurewitz from the Technion Biomedical Core Facility unit for assistance with the Automatic slide scanner and Olympus VS200 digital scanner – Brightfield imaging. We would like to thank Katren Sakran for the histology service.

## Author contributions

A.O. wrote the code and the manuscript. R.S. and M.A.S. provided support with image annotation and editorial work. I.L. and P.S., both pathologists, handled the muscle fibrosis analysis. A.O., A.S, and P.H. were responsible for conceptualization, supervision, funding acquisition, and reviewing, writing, and editing the manuscript.

## Competing interests

The authors declare no competing interests.

## Notes

https://github.com/Anas-Odeh/FibroTrack

### Competing Interest Statement

The authors have declared no competing interest.

## References

1. Bann, G. G. Antagonizing HFpEF by Targeting Fibrosis. Circulation 151, 396–399 (2025).

2. Odeh, A. et al. Anti-fibrotic, muscle-promoting antibody-drug conjugates for the improvement and treatment of DMD. iScience 28, 112335 (2025).

3. Mayorca-Guiliani, A. E. et al. ECM formation and degradation during fibrosis, repair, and regeneration. npj Metab Health Dis 3, 25 (2025).

4. Facchin, C. et al. FIBER-ML, an Open-Source Supervised Machine Learning Tool for Quantification of Fibrosis in Tissue Sections. Am J Pathol 192, 783–793 (2022).

5. Ségard, B.-D. et al. Quantification of fibrosis extend and airspace availability in lung: A semi-automatic ImageJ/Fiji toolbox. PLoS One 19, e0298015 (2024).

6. Sb, M., P, D., Bc, F. & P, C. Molecular imaging of fibrosis: recent advances and future directions. The Journal of clinical investigation 129, (2019).

7. Schipke, J. et al. Assessment of cardiac fibrosis: a morphometric method comparison for collagen quantification. J Appl Physiol (1985) 122, 1019–1030 (2017).

8. Y, H., et al. Image analysis of liver collagen using sirius red is more accurate and correlates better with serum fibrosis markers than trichrome. Liver international : official journal of the International Association for the Study of the Liver 33, (2013).

9. Hunter, R. S. Photoelectric Color Difference Meter*. J. Opt. Soc. Am. 48, 985 (1958).

10. Z, Y., et al. A deep learning-based approach for fully automated segmentation and quantitative analysis of muscle fibers in pig skeletal muscle. Meat science 213, (2024).

11. Mula, J., Lee, J. D., Liu, F., Yang, L. & Peterson, C. A. Automated image analysis of skeletal muscle fiber cross-sectional area. J Appl Physiol (1985) 114, 148–155 (2013).

12. Waisman, A., Norris, A. M., Elías Costa, M. & Kopinke, D. Automatic and unbiased segmentation and quantification of myofibers in skeletal muscle. Sci Rep 11, 11793 (2021).

13. Jocher, G., Qiu, J. & Chaurasia, A. Ultralytics YOLO. (2023).

14. Dwyer, B., Nelson, J. & Hansen, T. Version 1.0. Roboflow (2024).

15. Serrano, A. L. & Muñoz-Cánoves, P. Fibrosis development in early-onset muscular dystrophies: Mechanisms and translational implications. Semin Cell Dev Biol 64, 181–190 (2017).

16. Barnard, A. M. et al. Characterizing Expiratory Respiratory Muscle Degeneration in Duchenne Muscular Dystrophy Using MRI. Chest 161, 753–763 (2022).

17. Duca, F. et al. Cardiac extracellular matrix is associated with adverse outcome in patients with chronic heart failure. Eur J Heart Fail 19, 502–511 (2017).

