## Supplementary figures for "FibroTrack: A Standalone Deep Learning Platform for Automated Fibrosis Quantification in Muscle and Cardiac Histology"

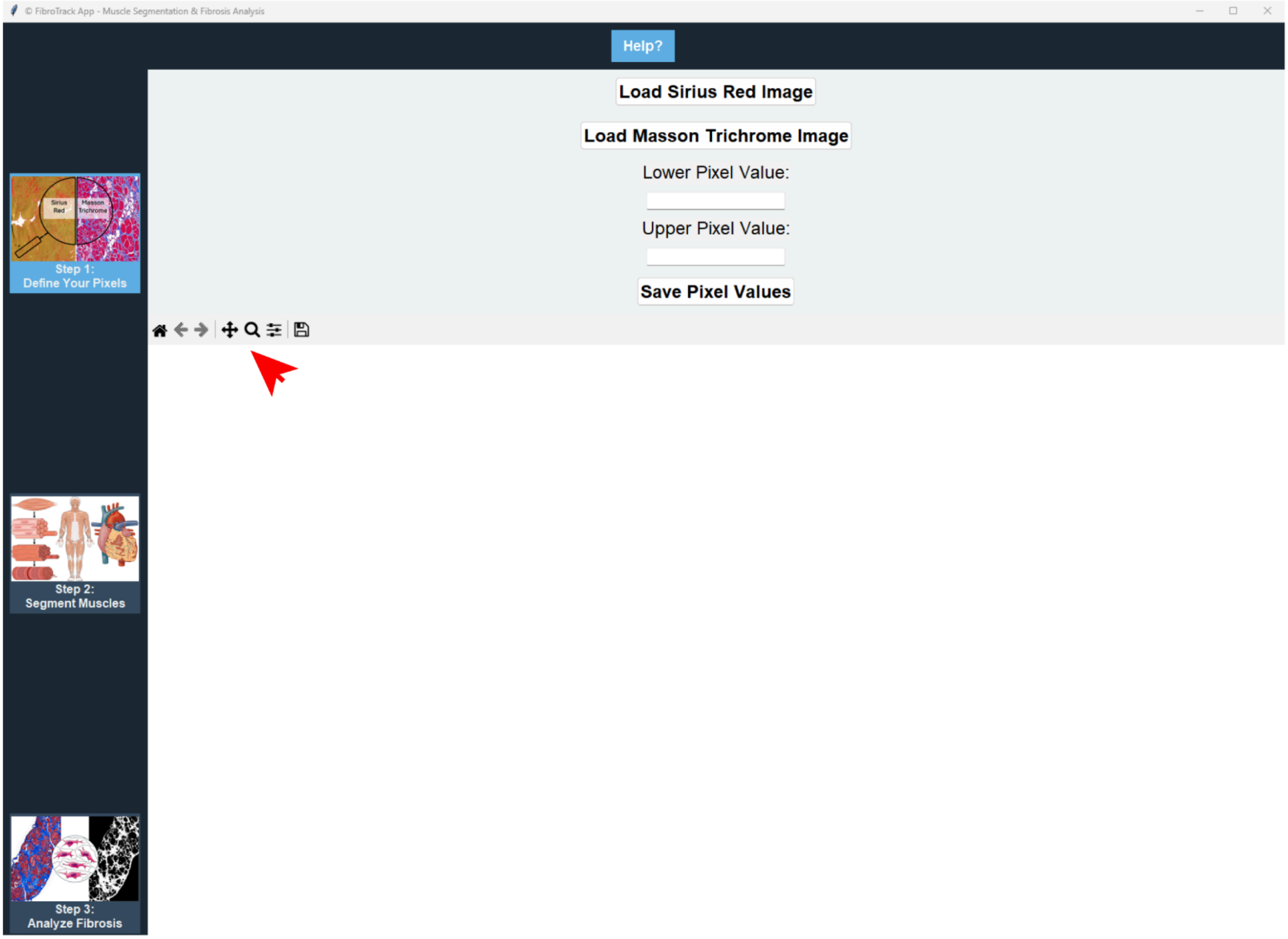


**Supplementary figure S1. Overview of the FibroTrack interface.**

A user interface for analyzing Sirius Red and Masson Trichrome stained histological images. The fibrotic region is segmented with the help of user-inputted lower- and upper-pixel intensity values that are defined based on an uploaded image. To make the user-friendly interface, tooltips were used, and it shows when hovering over buttons to guide what steps need to be done next in the process. This workflow is organized into the following three steps: (1) Define Your Pixels, (2) Segment Muscles, and (3) Analyze Fibrosis, under which each tool to perform specific analysis are provided in the left-hand navigation bar. The leftmost toolbar (indicated by red arrow) allows additional operations like zoom in and out, which are useful for a closer look of certain regions within the fibrotic muscles. Detailed explanations and step-by-step implementation for each of these stages are provided in subsequent figures.

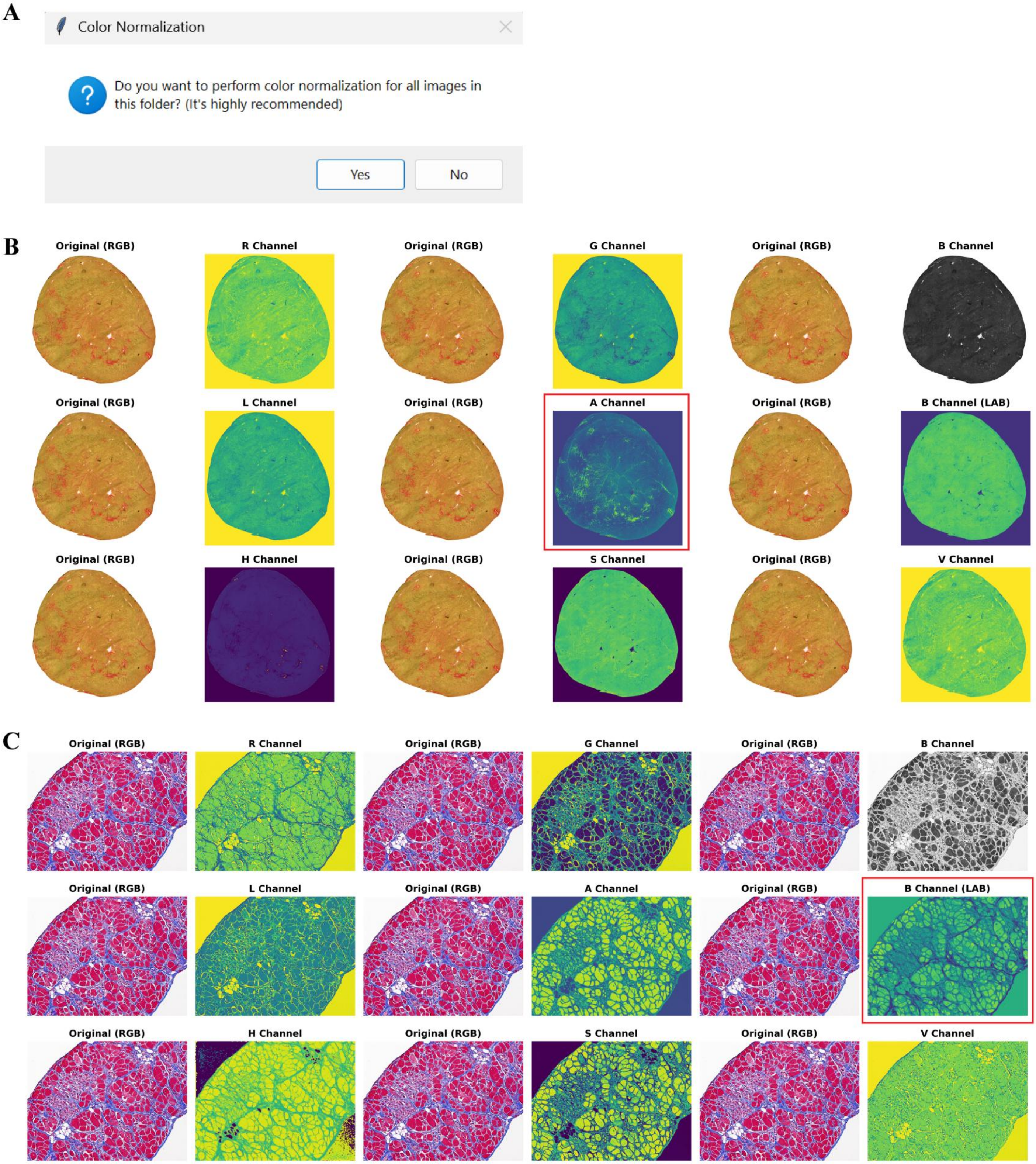


**Supplementary figure S2. Color normalization and optimal channel selection in FibroTrack.**

**(A)** Dialog box prompting users to perform color normalization on all images in the selected folder. **(B)** Comparative visualization of different color channels extracted from a Sirius Red-stained heart tissue. The original RGB image is shown alongside individual channel representations from RGB, HSV, and LAB color spaces. The A channel in LAB space (highlighted with red rectangle) provides optimal contrast for detecting collagen fibers stained with Sirius Red, as evidenced by the clear visualization of fibrotic structures in yellow-green against a darker background. **(C)** Similar channel comparison for a Masson's Trichrome-stained tissue sample. The B channel in LAB color space (highlighted with red rectangle) demonstrates superior ability to isolate fibrotic regions compared to other color spaces. In this representation, collagen fibers appear as distinct structures with enhanced contrast in purple against a brighter background, facilitating more accurate quantification of fibrosis.

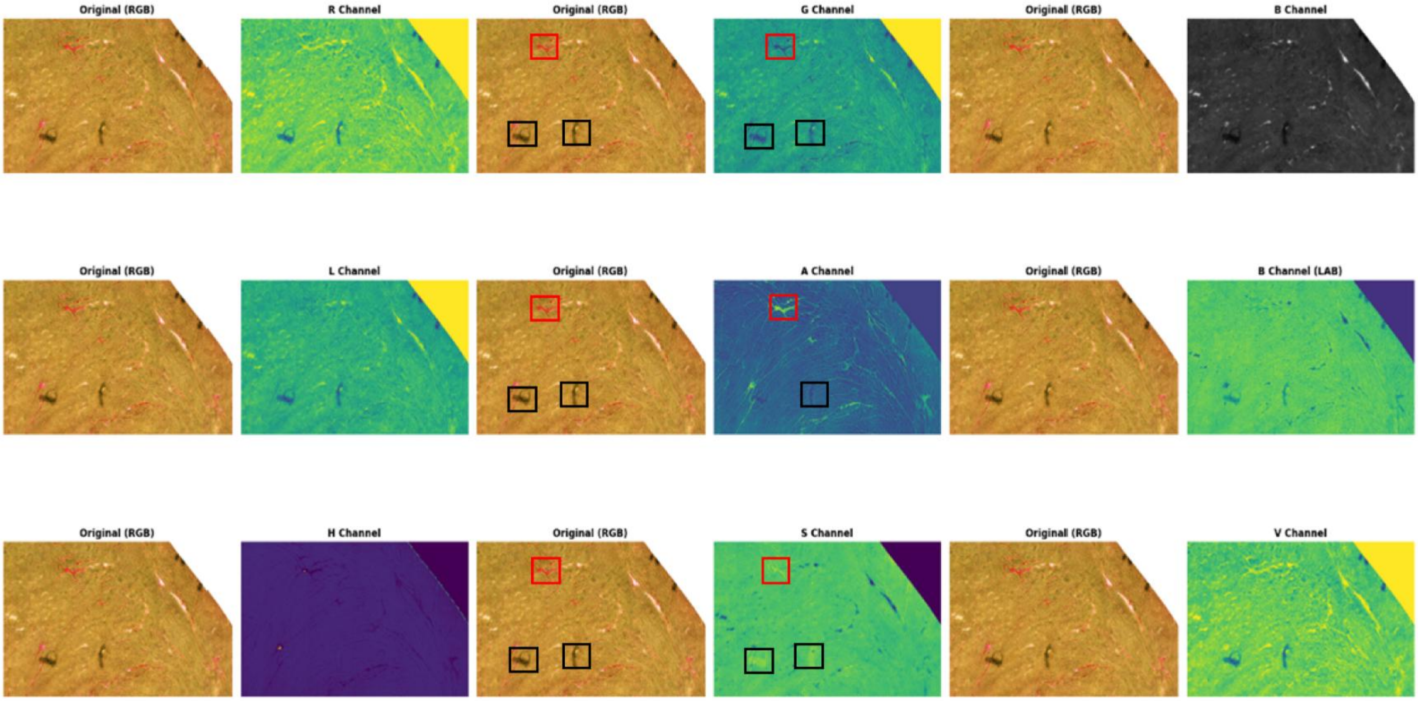


**Supplementary figure S3. Comparative Analysis of Color Space Channel Splitting in Cardiac Tissue (Sirius Red).**

Differential performance of RGB, LAB, and HSV color space channel splitting in Sirius Red-stained cardiac tissue samples. The upper panel displays three original RGB images with their corresponding individual color channels. In RGB splitting, while the G channel shows some discrimination of fibrotic areas (appearing as darker regions, indicated by red rectangle), it also captures non-fibrotic artifact staining (indicated by black rectangles). The middle panel displays LAB color space splitting, where the A channel provides superior separation of fibrotic tissue (red rectangle) with no detection of artifacts or non-fibrotic structures (black rectangle). The bottom panel shows HSV color space splitting, with multiple channels displaying inconsistent fibrosis separation and significant artifact inclusion. This comprehensive channel comparison demonstrates that the LAB color space, particularly its A channel, most accurately isolates fibrotic tissue from other cardiac tissue components and potential artifacts, making it the optimal choice for fibrosis quantification in Sirius Red-stained samples.

A

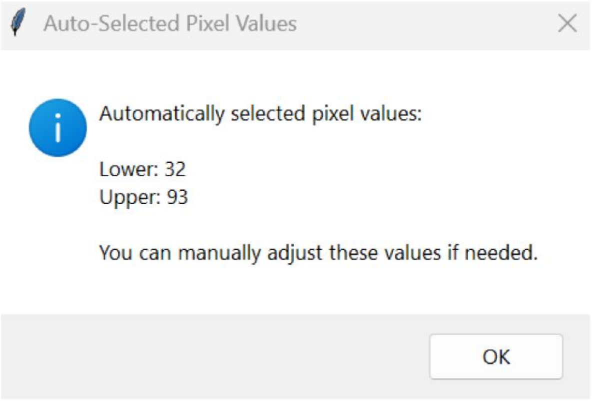


B

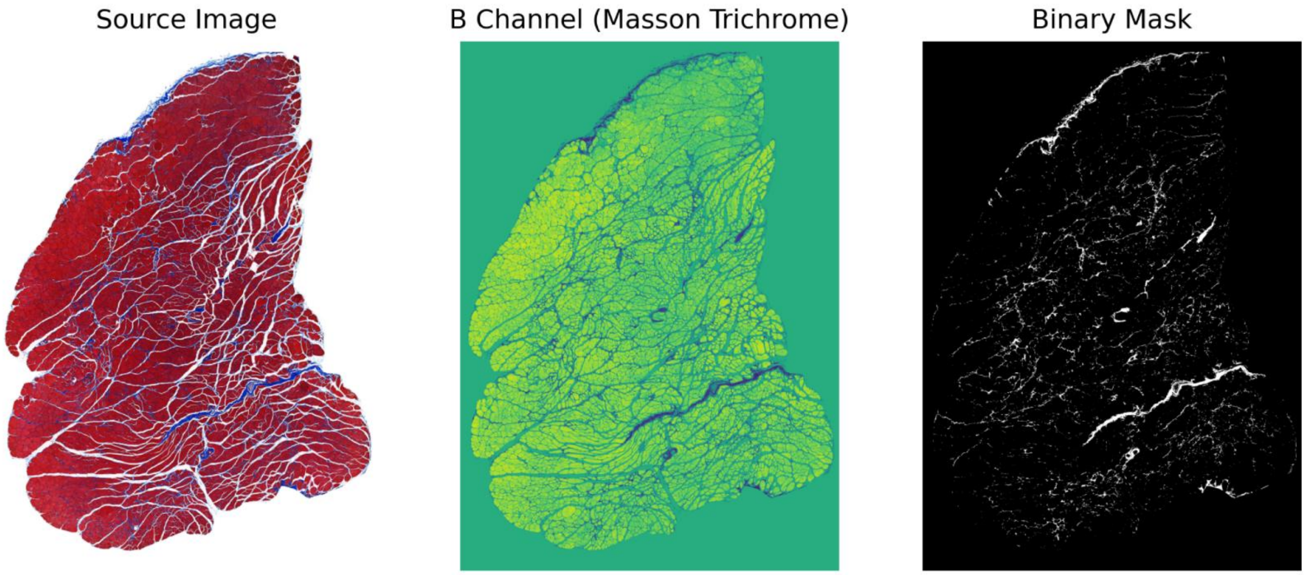


**Supplementary figure S4. Auto-selection of pixel values and result visualization in FibroTrack.**

**(A)** Pop-up window displaying automatically selected pixel intensity values for fibrosis detection. The software calculates optimal threshold values (Lower: 32, Upper: 93) based on LAB color space analysis, with a notification that users can manually adjust these values if needed. **(B)** Visualization of the fibrosis detection process in a Masson's Trichrome-stained tissue sample. Left panel shows the original source image with collagen fibers stained in blue; middle panel displays the B channel from LAB color space where fibrotic regions appear as lighter areas against a green background; right panel presents the resulting binary mask where white pixels represent detected fibrotic tissue based on the auto-selected pixel values.

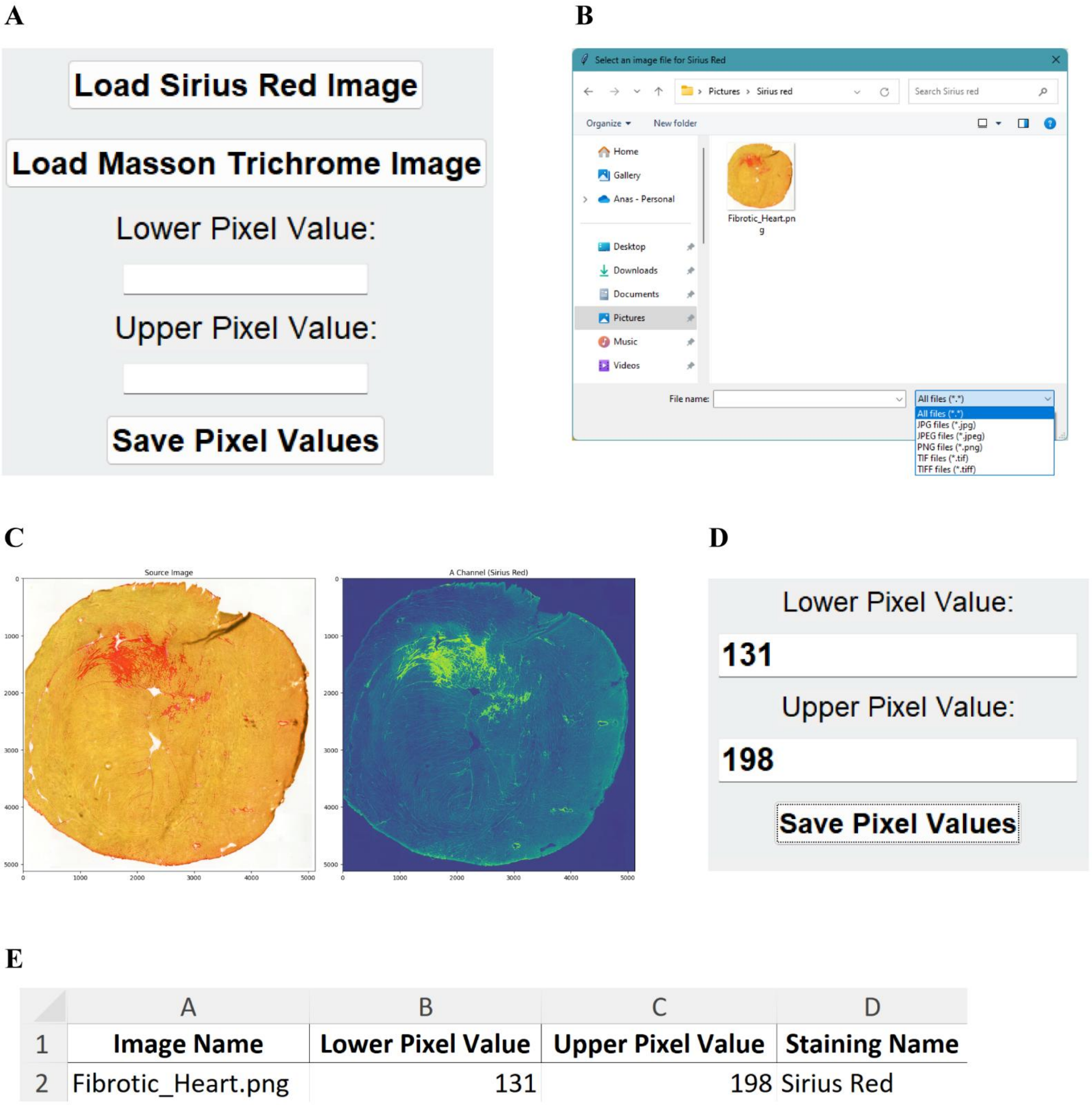


**Supplementary figure S5. Workflow for loading images and saving defined pixel intensity values.**

**(A)** FibroTrack interface for loading Sirius Red or Masson Trichrome images and defining the lower and upper pixel intensity values. **(B)** Selecting one image in one of the supported formats (jpg, jpeg, png, tif, tiff) to load into the interface. **(C)** The source image (left) and its LAB color space A channel (right) are displayed for pixel intensity value selection. **(D)** Defined lower- and upper-pixel intensity values are entered into the designated fields, and the "Save Pixel Values" button is clicked. **(E)** An Excel file containing the image name, lower pixel value, upper pixel value, and staining type (Sirius Red) is generated and saved.

**A**

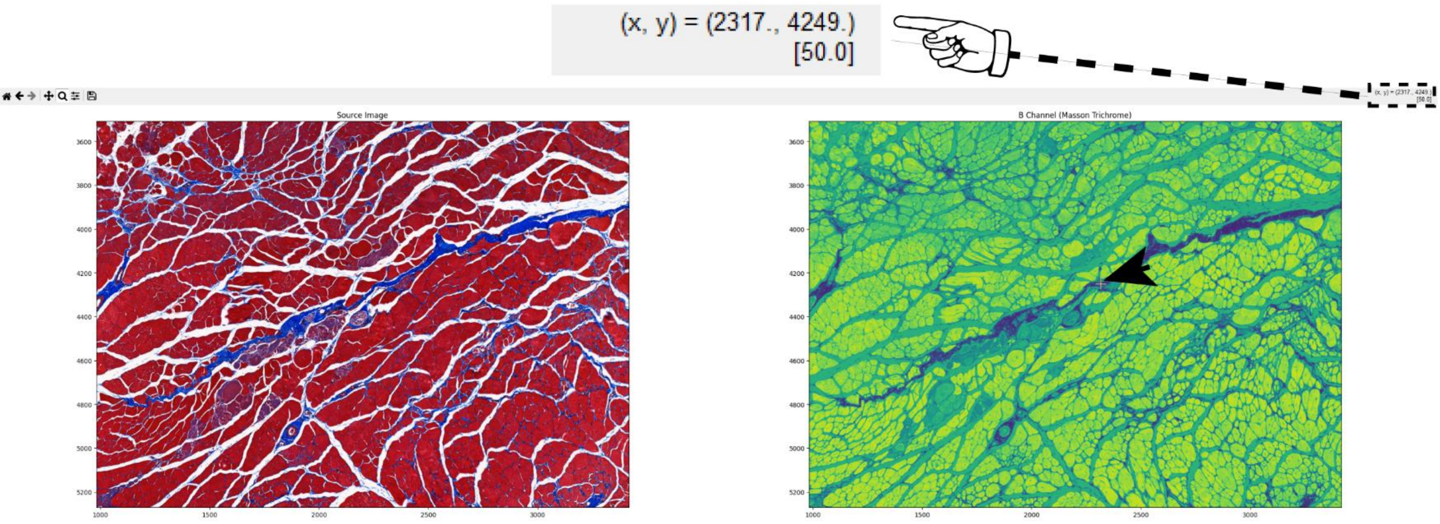


**B**

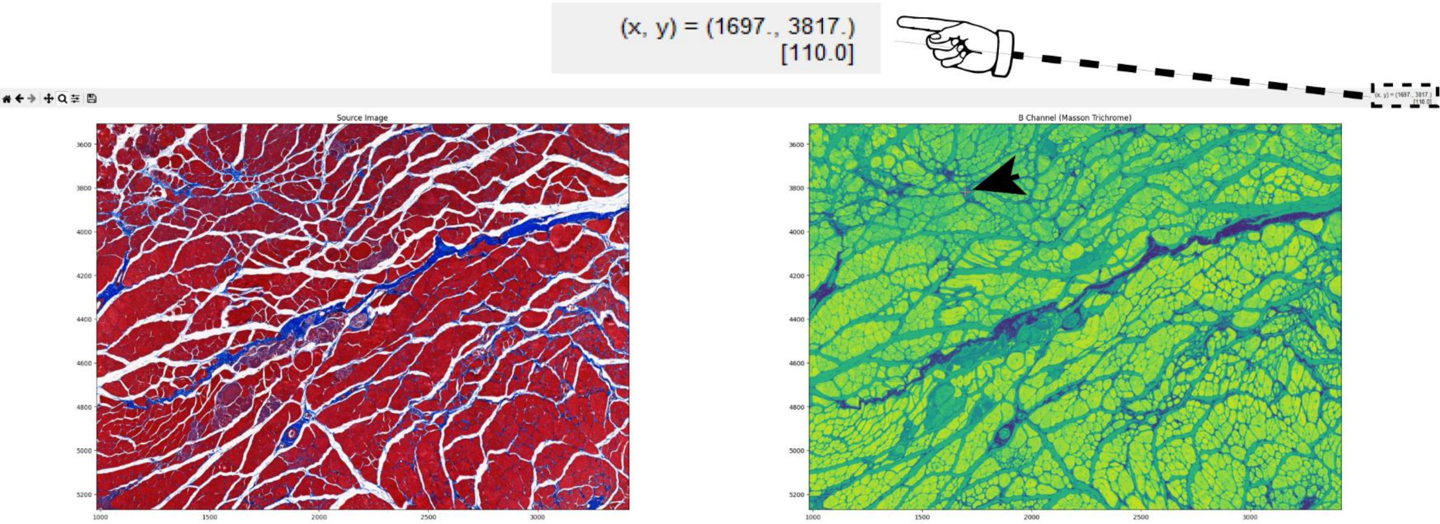


**Supplementary figure S6. Defining lower and upper pixel intensity values for fibrotic tissue in Masson Trichrome-Stained Images.**

**(A)** Defining of the lower pixel intensity value [50.0], showing the darkest fibrotic area in the B channel. The black arrow points to the picked pixel location in the B channel, and the coordinates (x, y = 2317, 4249) of the chosen pixel are shown in the interface. **(B)** The selection of the upper pixel intensity value [110.0], illustrating the brightest fibrotic area in the B channel. The black arrow highlights the selected pixel in the B channel, and the coordinates (x, y = 1697, 3817) of the selected pixel are displayed.

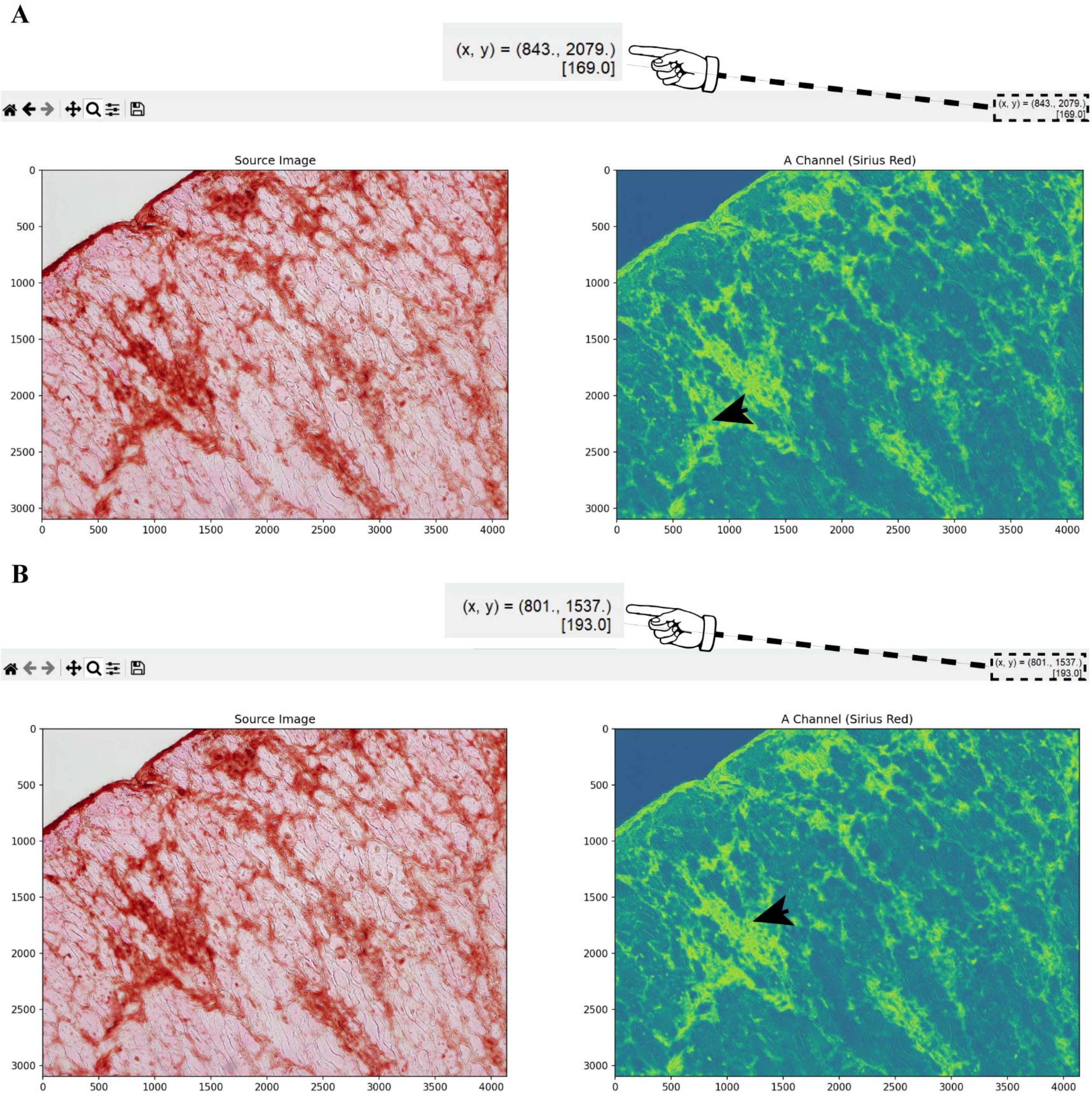


**Supplementary figure S7. Defining lower and upper pixel intensity values for fibrotic tissue in Immunohistochemistry for collagen I using the SR module.**

**(A)** The lower pixel intensity value was defined to [169.0], displaying the darkest pixel of fibrotic area. The black arrow indicates the pixel location selected whithin the image, which is indicated in the interface by its pixel coordinates (x, y = 843, 2079). **(B)** The highest pixel intensity value selected [193.0], showing the brightest fibrotic area. The coordinates of the selected pixel (x, y = 801, 1537) is indicated by the black arrow.

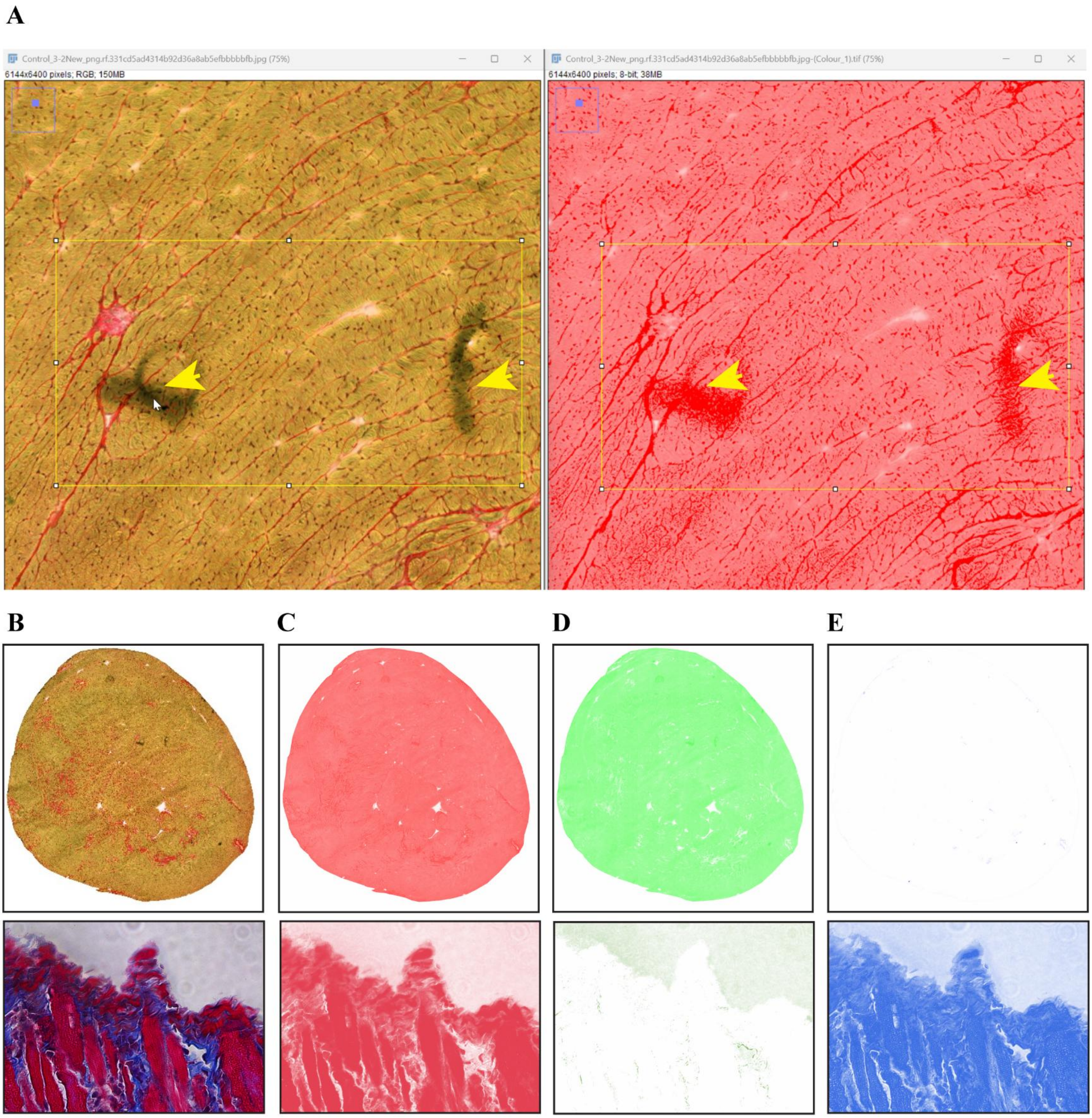


**Supplementary figure S8. Comparison of Manual Thresholding and Color Deconvolution Techniques for Fibrosis Analysis.**

**(A)** Heart tissue images shown in ImageJ. Source RGB image (left) and its RGB-color-deconvolution image, red channel (right). The original stained tissue clearly shows artifacts (black spots indicated by yellow arrows) that could be misinterpreted as fibrotic regions, while the deconvolution highlights all red-stained areas indiscriminately, failing to distinguish between true fibrosis and artifacts. **(B-E)** Comparative analysis of different color processing methods for cardiac tissue samples. **(B)** Original RGB source images of cardiac tissue samples stained with both Sirius Red (top panel) and Masson Trichrome (bottom panel). **(C-E)** Individual color channels after deconvolution: **(C)** Red channel,

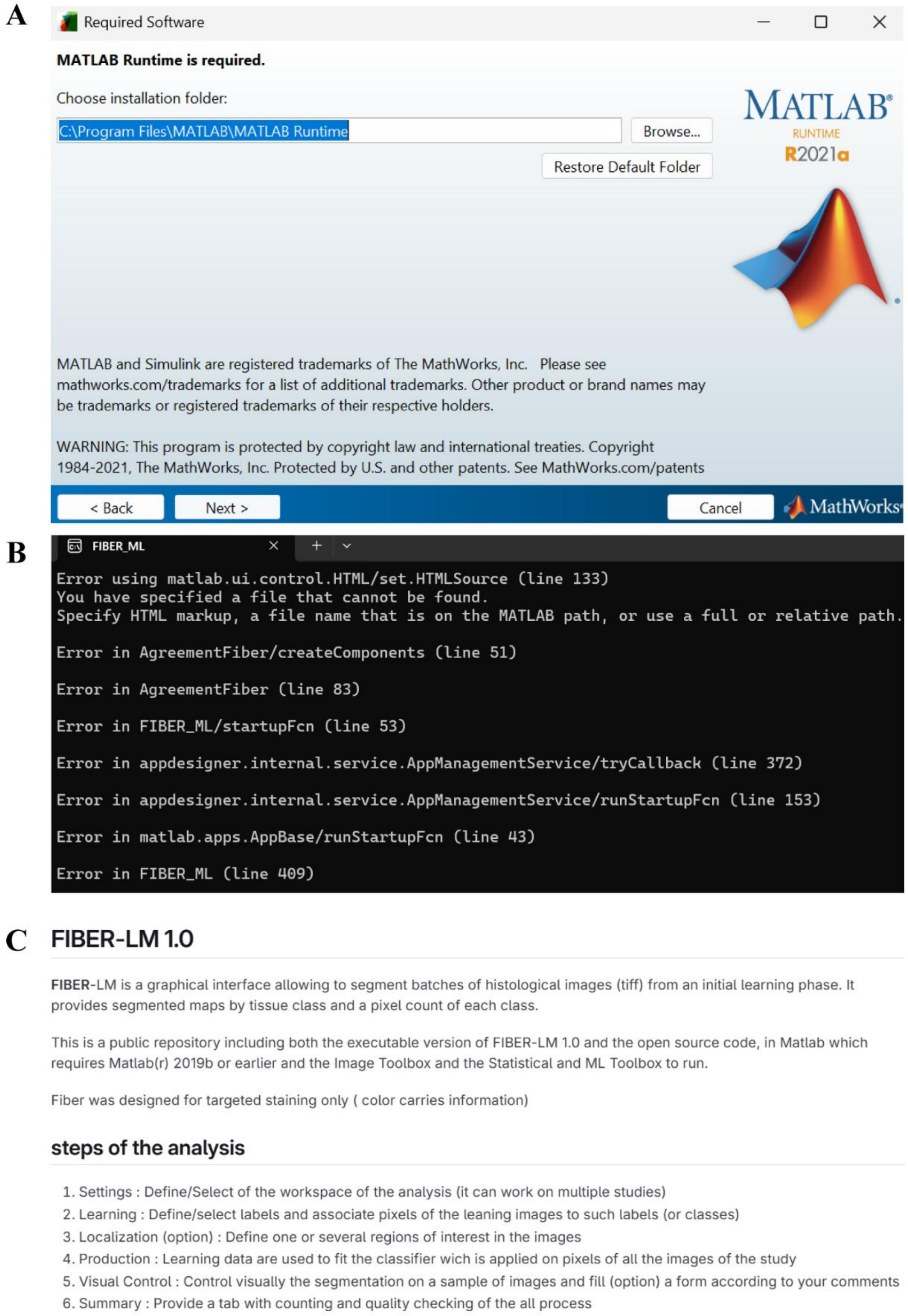


**Supplementary figure S9. Implementation Barriers of FIBER-ML Software.**

**(A)** MATLAB Runtime installation window for FIBER-ML, showing the first of several required software dependencies. **(B)** Command prompt displaying multiple execution failures, demonstrating how even users with programming expertise encounter technical issues when attempting to run the software. **(C)** FIBER-ML documentation revealing additional requirements including specific MATLAB version (v2019b or earlier) plus specialized toolboxes, along with a complex multi-step manual workflow.

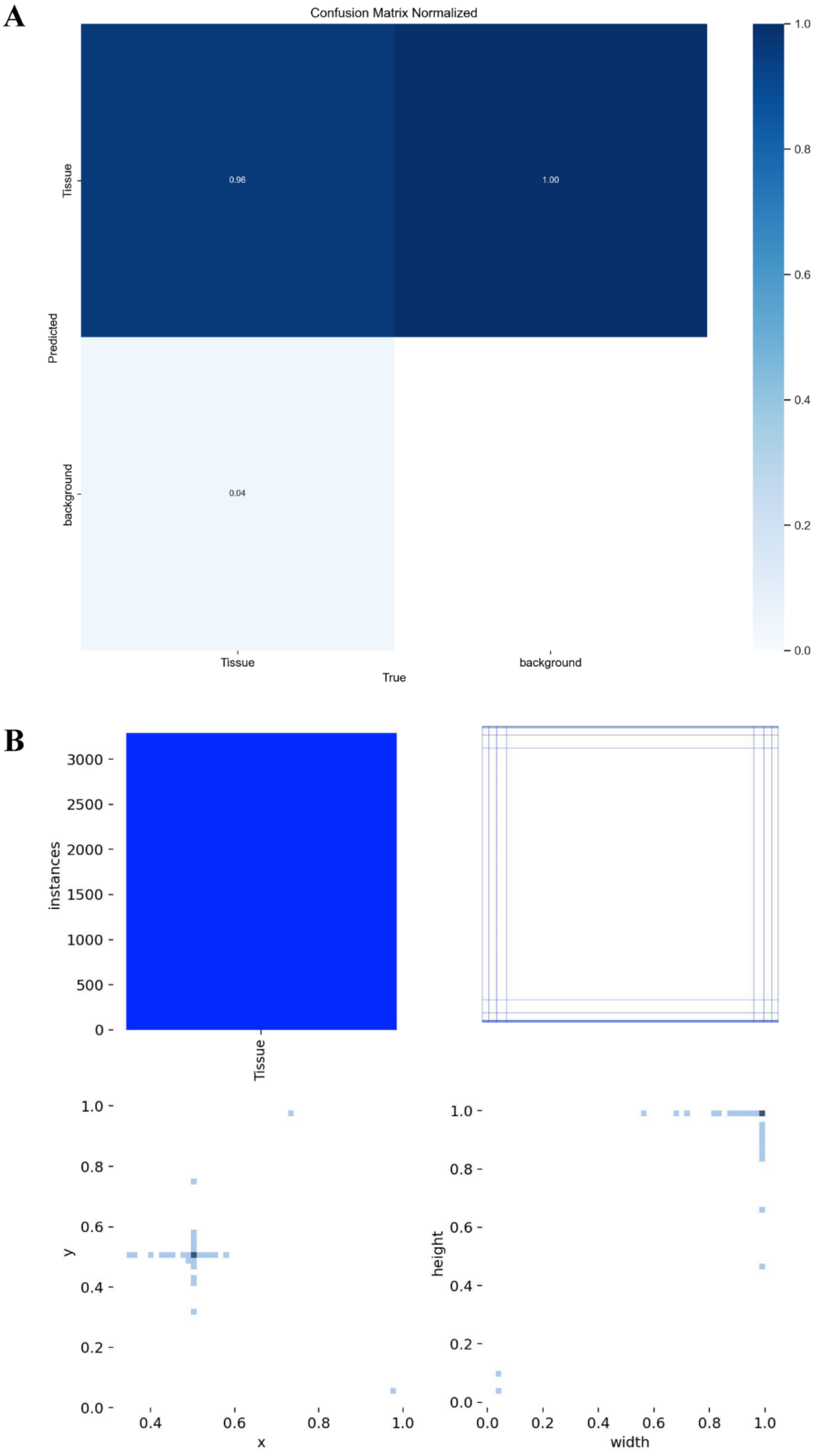


**Supplementary figure S10. Performance and error distribution of FibroTrack's tissue segmentation.**

**(A)** Confusion matrix showing segmentation performance across the test dataset (n=406): 96% sensitivity (true positive rate) for tissue pixels, 100% specificity (true negative rate) for background, 4% false negative rate, and 0% false positive rate. The model achieved an F1-score of 97%, demonstrating excellent balance between precision and recall in tissue segmentation. **(B)** Spatial analysis of segmentation performance. Heatmaps (top) and scatter plots (bottom) demonstrate consistent segmentation accuracy across spatial coordinates and tissue dimensions. The uniform distribution of high-performance metrics across all positions (x-y coordinates) and size parameters (width-height) confirms that FibroTrack maintains reliable segmentation regardless of tissue morphology or positioning within the image frame.

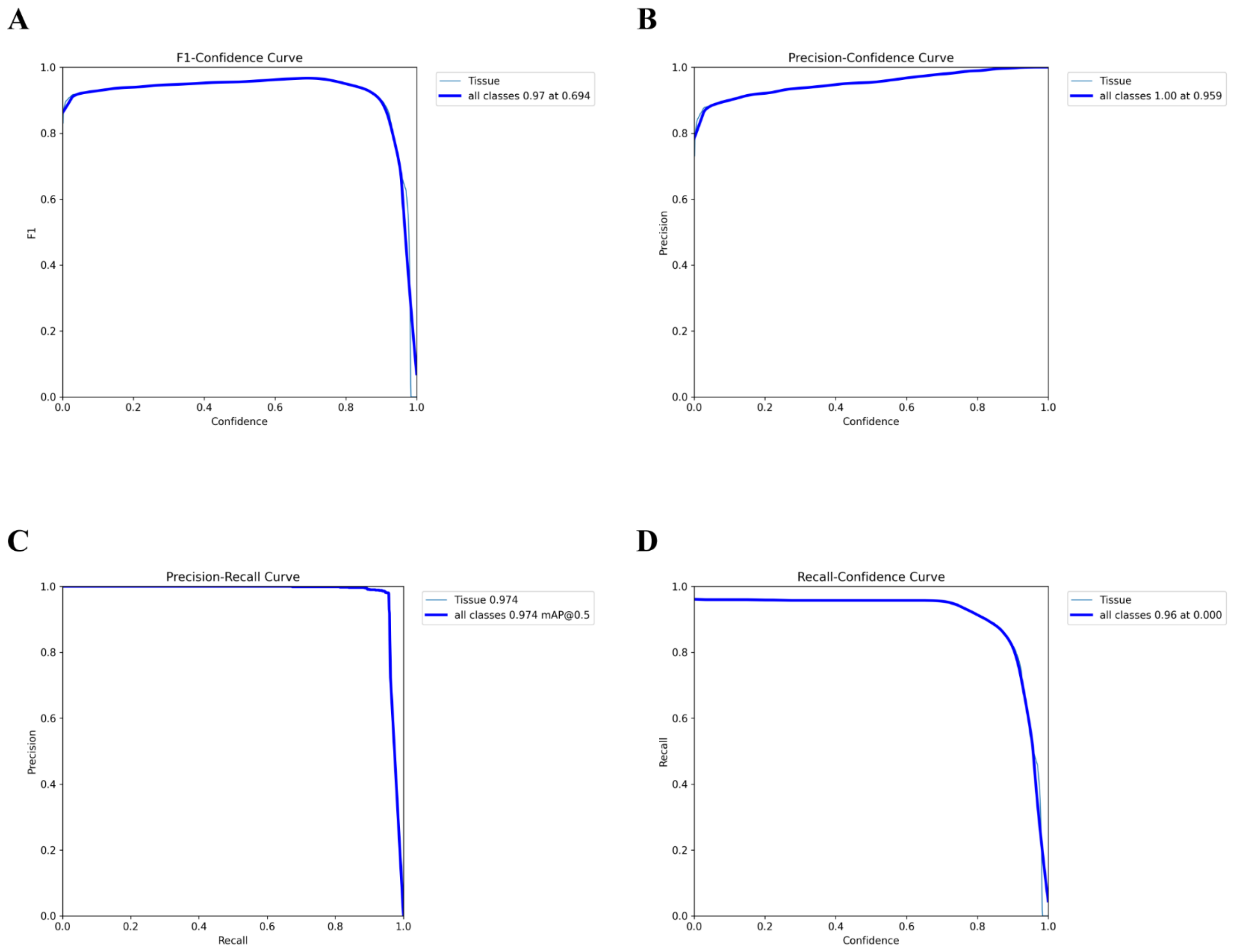


**Supplementary figure S11. Bounding box confidence curves for model evaluation.**

**(A)** F1-Confidence Curve shows the F1 score as a function of confidence. The model achieves a high F1 score (0.97) at a confidence threshold of (0.694). **(B)** Precision-Confidence Curve represents the precision as a function of confidence. The model achieves a high precision (1.00) at a confidence threshold of (0.959). **(C)** Precision-Rcall Curve displays the relationship between precision and recall, with the model achieving an mAP@0.5 of (0.974). **(D)** Recall-Confidence Curve shows recall as a function of confidence. The model achieves a recall of (0.96) at a confidence threshold of (0.000).

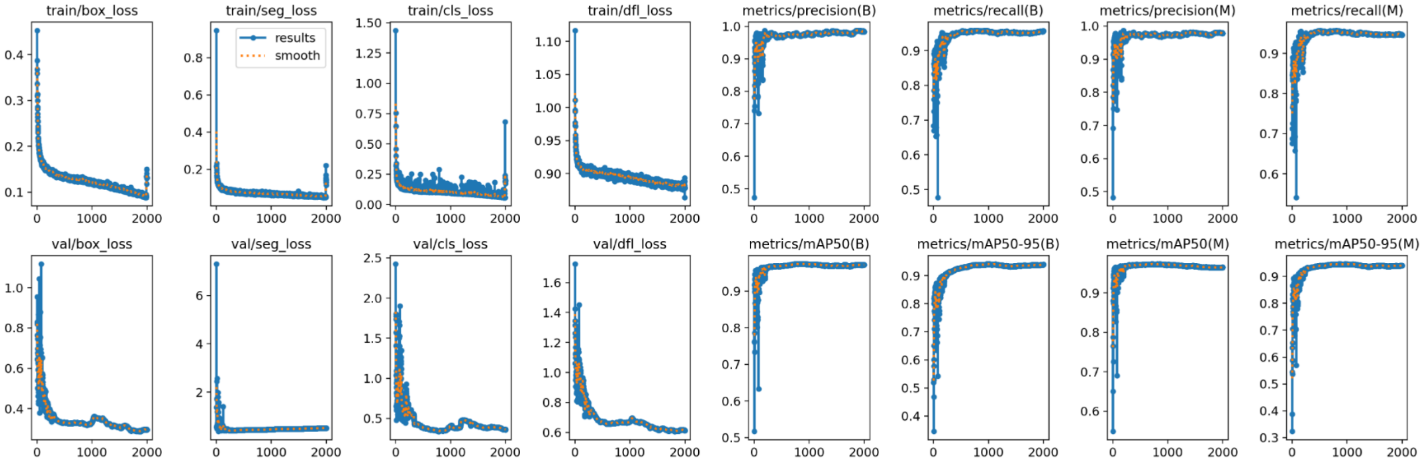


**Supplementary figure S12. Training and validation loss and metrics.**

This figure shows 16 plots tracking the YOLOv11 model's training and validation performance over time. The first four plots on the top left row display training losses for bounding box predictions, segmentation, classification, and distribution focal loss (DFL), with decreasing trends indicating improved model learning. The next four plots show precision and recall for both bounding boxes (B) and masks (M), with high values reflecting accurate predictions. The bottom row shows corresponding validation losses and mean average precision (mAP) at IoU thresholds of 0.5 and 0.5-0.95, highlighting the model’s generalization and detection accuracy.

### Table S1.

| Class Name | Images | Instances | Box Precision (P) | Box Recall (R) | Box $mAP_{(50-95)}$ | Mask Precision (P) | Mask Recall (R) | Mask $mAP_{50}$ | Mask $mAP_{(50-95)}$ |
|---|---|---|---|---|---|---|---|---|---|
| Cardiac and skeletal muscles | 905 | 937 | 0.956 | 0.974 | 0.941 | 0.975 | 0.953 | 0.972 | 0.946 |

**Supplementary Table S1. Validation Metrics for Cardiac and Skeletal Muscle Class.**

The validation metrics for the cardiac and skeletal muscle class with YOLOv11n-seg model. Results on 905 images with a total of 937 instances. The data features low variance and the model's performance shows high precision (P) and recall (R), for both bounding boxes as well as segmentation masks. The bounding box precision and recall are 0.956 (95.6%), 0.974 (97.4%) respectively with a box (mAP50-95) of 0.941. The results for the segmentation masks are a P=0.975, R=0.953 (95.3%) and mask (mAP50-95) of 0.946 (94.6%). This displays strong detection and segmentation for both cardiac as well as skeletal muscle tissues.

**Table S2.**

| Image Name | area | centroid-0 | centroid-1 | orientation | eccentricity | extent | perimeter | solidity | axis major length | axis minor length |
|---|---|---|---|---|---|---|---|---|---|---|
| mdx1 | 2268370 | 769.97 | 1028.73 | 1.21 | 0.87 | 0.72 | 6866.38 | 0.95 | 2486.27 | 1217.14 |
| mdx2 | 2584061 | 750.41 | 1206.24 | 1.13 | 0.56 | 0.88 | 6879.56 | 0.98 | 2031.26 | 1686.91 |
| mdx3 | 2610510 | 751.74 | 1166.71 | 1.10 | 0.61 | 0.86 | 7057.87 | 0.97 | 2091.31 | 1657.87 |
| mdx4 | 2710959 | 793.85 | 1066.89 | 1.19 | 0.60 | 0.87 | 7149.30 | 0.95 | 2116.93 | 1701.17 |
| mdx5 | 2752732 | 766.26 | 1053.92 | 1.14 | 0.63 | 0.86 | 7363.15 | 0.96 | 2180.58 | 1691.80 |
| mdx6 | 2567446 | 778.62 | 1028.86 | 1.28 | 0.81 | 0.82 | 7012.68 | 0.89 | 2412.64 | 1402.89 |
| mdx7 | 11396704 | 1639.84 | 2233.41 | -1.27 | 0.65 | 0.91 | 13773.44 | 0.99 | 4475.40 | 3393.12 |
| mdx8 | 10790921 | 1692.31 | 2332.25 | -1.16 | 0.70 | 0.85 | 13602.70 | 0.99 | 4501.18 | 3235.57 |
| mdx9 | 12149820 | 1613.43 | 2152.13 | -1.41 | 0.68 | 0.95 | 14010.05 | 1.00 | 4679.96 | 3425.83 |
| mdx10 | 307618 | 1408.51 | 1228.23 | 1.54 | 1.00 | 0.66 | 5198.94 | 0.73 | 2392.23 | 217.11 |
| mdx11 | 1012863 | 1280.84 | 1097.70 | 1.55 | 0.96 | 0.96 | 4973.85 | 1.00 | 2251.87 | 596.84 |

**Supplementary Table S2. Quantitative Muscle Region Properties from FibroTrack Segmentation.**

Table summarizing key quantitative properties for muscle region segmentation. Metrics include geometric (Area, Perimeter, Axis Lengths), positional (Centroid Coordinates), and shape descriptors (Orientation, Eccentricity, Extent, Solidity), highlighting the detailed output generated by FibroTrack's image segmentation process.
